# Climate Impacts on Sockeye Salmon Productivity Vary Across Life Stages and Regions

**DOI:** 10.64898/2026.08.26.746844

**Authors:** Jan F. Finke, Travis C. Tai, Cameron Freshwater, Brendan Connors, Amber M. Holdsworth, Greig L. Oldford, Daniel T. Selbie, Howard W. Stiff, Patrick L. Thompson

**Affiliations:** Pacific Science Enterprise Centre, Fisheries and Oceans Canada, West Vancouver, BC, Canada; Pacific Biological Station, Fisheries and Oceans Canada, Nanaimo, BC, Canada; Institute of Ocean Sciences, Fisheries and Oceans Canada, Sidney, BC, Canada; Cultus Lake Salmon Research Laboratory, Fisheries and Oceans Canada, Cultus Lake, BC, Canada

**Keywords:** Keywords: Sockeye Salmon, Lifecycle, Climate, Causal Inference, Freshwater, Marine

## Abstract

Many Sockeye salmon (*Oncorhynchus nerka*) populations have declined over recent decades, and climate change is likely to exacerbate these declines through direct and indirect ecological effects. The response to the associated environmental changes is likely to vary among life stages, populations, and regions. Quantitative estimates of climate change driven impacts that account for this variability could fill a critical gap and provide forward-looking insights into how sockeye are expected to respond to future climate-driven change across their lifecycle. To address this need we developed a hierarchical population dynamics model parameterized with juvenile, adult return and spawner abundance data from 13 sockeye salmon populations from Washington State to northern British Columbia. We used a formal causal inference framework that paired salmon abundance data with a suite of environmental covariates hypothesized to represent ecological conditions across the lifecycle. We used the model to estimate population-specific responses to each environmental driver, then combined parameter estimates with projections from down-scaled climate change models to estimate productivity responses to anticipated environmental change. We found that historical sockeye productivity was strongly associated with environmental covariates, which explained more interannual variability in return abundance than spawner abundance in most populations. However, the life stages and specific environmental covariates with the largest impacts differed among populations and regions, often displaying a latitudinal gradient. Increases in coastal ocean temperatures and mixed layer depth generally had negative effects though they varied among regions. Increased freshwater summer rearing and return migration temperatures had weaker but consistently negative effects. Under future climate conditions, projected changes in these environmental covariates are expected to result in substantial declines in productivity across most populations. Sockeye salmon display varying degrees of sensitivity to climate change across life stages, populations, and regions. Effective future management will require explicitly accounting for these life stage and population-specific responses.

## 1 Introduction

Many populations of anadromous Pacific salmon (*Oncorhynchus spp.*) have declined over recent decades despite large reductions in harvest (Grant 2019, Walters et al. 2019). Changes in temperature, oceanographic conditions and freshwater flow, and related ecosystem changes are all hypothesized to contribute to these trends. These environmental changes are expected to accelerate over the coming decades (Poertner et al. 2022), and will likely impact salmon throughout freshwater and marine life stages. However, despite an increasing availability of climate model projections, quantitative estimates of climate-driven impacts on salmon are limited, particularly regarding how effects vary among life stages and populations. This information is needed to support the long term management strategies required to sustain salmon into the future.

Sockeye salmon (*Oncorhynchus nerka*) utilize diverse freshwater and marine habitats and hence experience environmental change across disparate ecosystems, each posing unique challenges (Healey 2011, Crozier et al. 2021). Rearing in freshwater streams and lakes, sockeye salmon are exposed to warming waters, as well as changing precipitation and flow rates, which impact them directly and indirectly through changes in food-web dynamics (Carter et al. 2017). Once in the ocean, sockeye salmon join a complex food web that includes their prey, competitors, and predators, all of which are responding to changes in temperature and ocean conditions (Tucker et al. 2015, Farley et al. 2007, McKinnell and Irvine 2021, Connors et al. 2025). Furthermore, the marine environments they traverse are far from homogeneous; as sockeye salmon migrate along the coastal continental shelf to the open North Pacific, they encounter oceanographic domains with distinct physical characteristics and ecological communities. Migrating up-river to their spawning grounds, sockeye salmon are exposed to varying flow rates and warming water temperatures that are increasingly placing them under physiological stress (Eliason et al. 2011, Martins et al. 2011, Crozier et al. 2020, Atlas et al. 2021).

Correlated environmental covariates and unobserved processes can bias parameter estimates, resulting in misleading conclusions about how ecosystems respond to changing conditions (McElreath 2020). For example, regional climate forcing can cause environmental conditions in different habitats to covary (Mantua and Hare 2002, Gosselin et al. 2021), making it difficult to distinguish the degree to which each is responsible for observed changes in salmon survival. Additionally, when key variables such as prey abundance are unobserved, it is important to recognize that the estimated effect of environmental covariates like temperature reflect both their direct physiological impacts and their indirect influence through the food-web (Cinelli et al. 2024). Careful model development and interpretation grounded in causal-inference principles can be used to mitigate these risks (Pearl 2009, Correia et al. 2026, Cham-pagnat et al. 2026). Framing model relationships causally allows researchers to express assumptions clearly, explicitly define what model coefficients represent, and guard against misinterpreting correlated environmental signals. These causal frameworks are particularly important when trying to use historical relationships among productivity and covariates to predict future responses under climate change.

Past approaches to assess climate change impacts on Pacific salmon, provided only a partial view of how impacts vary across life stages and regions. Qualitative vulnerability assessments identify populations most vulnerable to climate change based on expert evaluation of biological sensitivity, exposure, and adaptive capacity, but they do not quantify links between environmental conditions and survival (Crozier et al. 2019). Quantitative hierarchical models relating total lifecycle productivity and environmental covariates have highlighted regional variability in the sensitivity of populations to covariates such as sea surface temperature, but provide limited insight into life stage specific survival, and may be confounded by ecosystems process not considered (Connors et al. 2020). Finally, quantitative lifecycle models applied within individual watersheds or with limited covariates have identified particularly vulnerable life stages and significant covariates under anticipated environmental change (Cun- ningham et al. 2018, Atlas et al. 2021, Akbarzadeh et al. 2021, Crozier et al. 2021), but their single-system or single-variable focus limit broader comparisons. Consequently, a cross-regional, life stage explicit, and causal inference-based framework that quantifies how freshwater and marine conditions influence survival is still lacking.

This project paired a network of sockeye salmon populations with environmental variables to quantify life stage specific responses and estimate sensitivity to future environmental change. We developed a hierarchical, lifecycle model that integrates juvenile, adult return, and adult spawner abundances with freshwater and marine environmental variables in a causal inference framework (detailed in Supplementary Section 5.1, Figure S2). The analysis spanned populations across a broad latitudinal gradient, from the Columbia River to the trans-boundary region in northern British Columbia. Our model estimated freshwater and marine productivity as functions of environmental conditions, while accounting for variable age structure and hatchery enhancement. We incorporated indices of freshwater and marine temperature, coastal mixed layer depth, and river discharge as key physical drivers that are hypothesized to impact productivity. Using these estimated relationships, we quantified changes in total lifecycle productivity due to projected climate change. Our analysis identified life history stages that have a disproportionate impact on productivity under changing climate conditions, providing a basis for tailored interventions across populations.

## 2 Methods

### 2.1 Salmon Lifecycle Data and Study Area

We fit a hierarchical model to data from lake-rearing sockeye salmon populations that span a wide geographical range from Washington State (USA) to the transboundary region of northern British Columbia (Canada) and southeast Alaska (USA). We grouped populations into three regional domains with distinct oceanographic properties along a latitudinal gradient. Specifically, the dataset is comprised of 13 populations that are part of the Columbia River (2) and Somass River (2) in the southern domain, the Fraser River (6) in the Fraser domain, and one population each from the Skeena River, Stikine River and Taku River in the northern domain (Figure 1). The populations from the Columbia and Somass systems were grouped into one domain because they both enter marine waters influenced by the California current. In addition, the small number of populations in each watershed (n = 2) was insufficient to estimate distinct hierarchical structure. Although freshwater conditions in the Somass and Columbia differ substantially, we retained the hierarchical structure in the analysis, with the ex- pectation that there may be high among population variability in freshwater responses in this domain.

**Figure 1:**
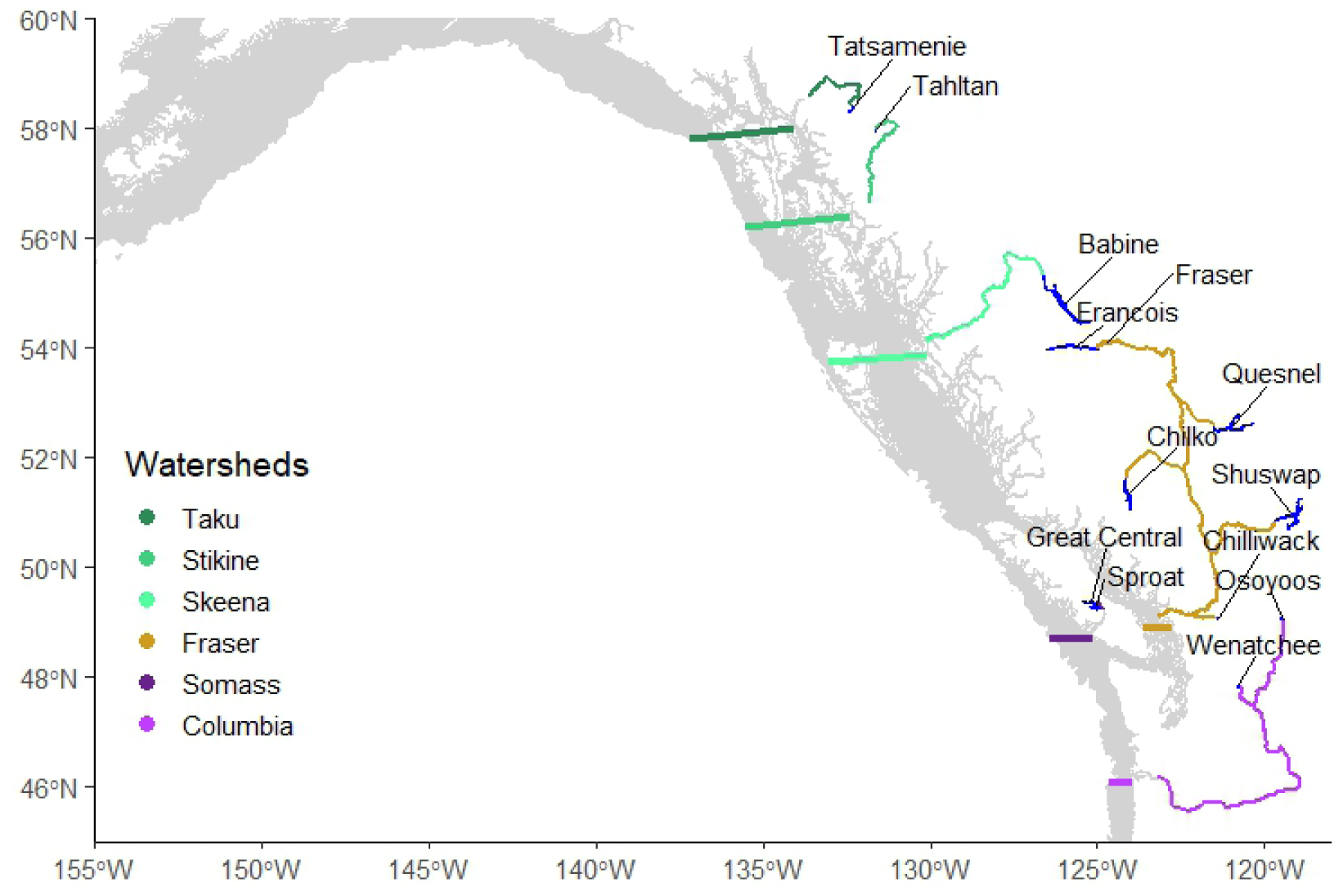
Populations with their associated rearing lakes, freshwater migration routes, ocean entry points and coastal migration routes. Horizontal lines indicate the extent of southern borders for ocean entry regions during coastal migration, colour coding indicates the domains we considered in the analysis, including Southern (purple), Fraser (gold), and Northern (green) regions. Grey shading indicates the extent of the coastal shelf down to 500 meters depth.

Observed abundance data included estimates of spawners (escapement), and juveniles (either fry or smolt stages). We paired this with year-specific observations of age composition within each life stage. For juveniles, age classes reflect the number of freshwater winters prior to smolting. For spawners, it is the combination of freshwater and marine winters experienced. We calculated adult return abundance (i.e., adult recruits) as a function of harvest rate and spawner abundance. We did not incorporate en route mortality rate estimates because it was unavailable for most populations (additional details below). For populations with hatchery supplementation (Osoyoos, Wenatchee, Tatsamenie, Tahltan), hatchery-origin fish were excluded when estimating natural-origin production of returns in the previous generation, but were included in post-harvest spawner abundance to quantify contributions to the next generation. The combined data spanned 44 years from 1981-2024; missing abundance, harvest, and/or age composition data were imputed implicitly within the Bayesian joint-likelihood model to propagate uncertainties through the model.

### 2.2 Environmental Covariates and Parameters

We incorporated a suite of freshwater and marine environmental covariates from different sources (Figure S1) to represent ecosystem conditions experienced by salmon during juvenile freshwater rearing, early marine entry, offshore marine residence, and upstream spawning migration. Freshwater productivity is affected by summer and winter rearing temperatures, smolt to recruit survival is affected by marine entry temperature and mixed layer depth (MLD), summer and winter open ocean temperatures, and return migration temperature and discharge, spawner abundance is affected by harvest and hatcheries (Figure 2). Covariates were assembled as annual time series and, depending on life history stage, covariates were either population-specific (fresh-water covariates), shared among populations within a watershed that have common marine migration corridors (early marine covariates), or shared among all populations (offshore marine covariates). Missing observations were imputed implicitly within the Bayesian joint-likelihood model, ensuring that uncertainty in environmental covariates propagated through to the estimates of life-history parameters. Environmental covariates were aggregated to spatial and temporal scales specific to each population, based on known or hypothesized freshwater and marine habitat use and life-history timing. For each population, covariates were summarized over relevant freshwater rearing areas, associated river mouth-coastal regions, and the open-ocean region, and temporally aligned to biologically meaningful periods including egg incubation, juvenile rearing, smolt out-migration, and marine residence (Figure S4). This approach ensured that environmental variations reflected conditions hypothesized to be experienced by each population at critical life-history stages. To allow for comparisons of their relative effects all covariates were centred and scaled as detailed below.

**Figure 2:**
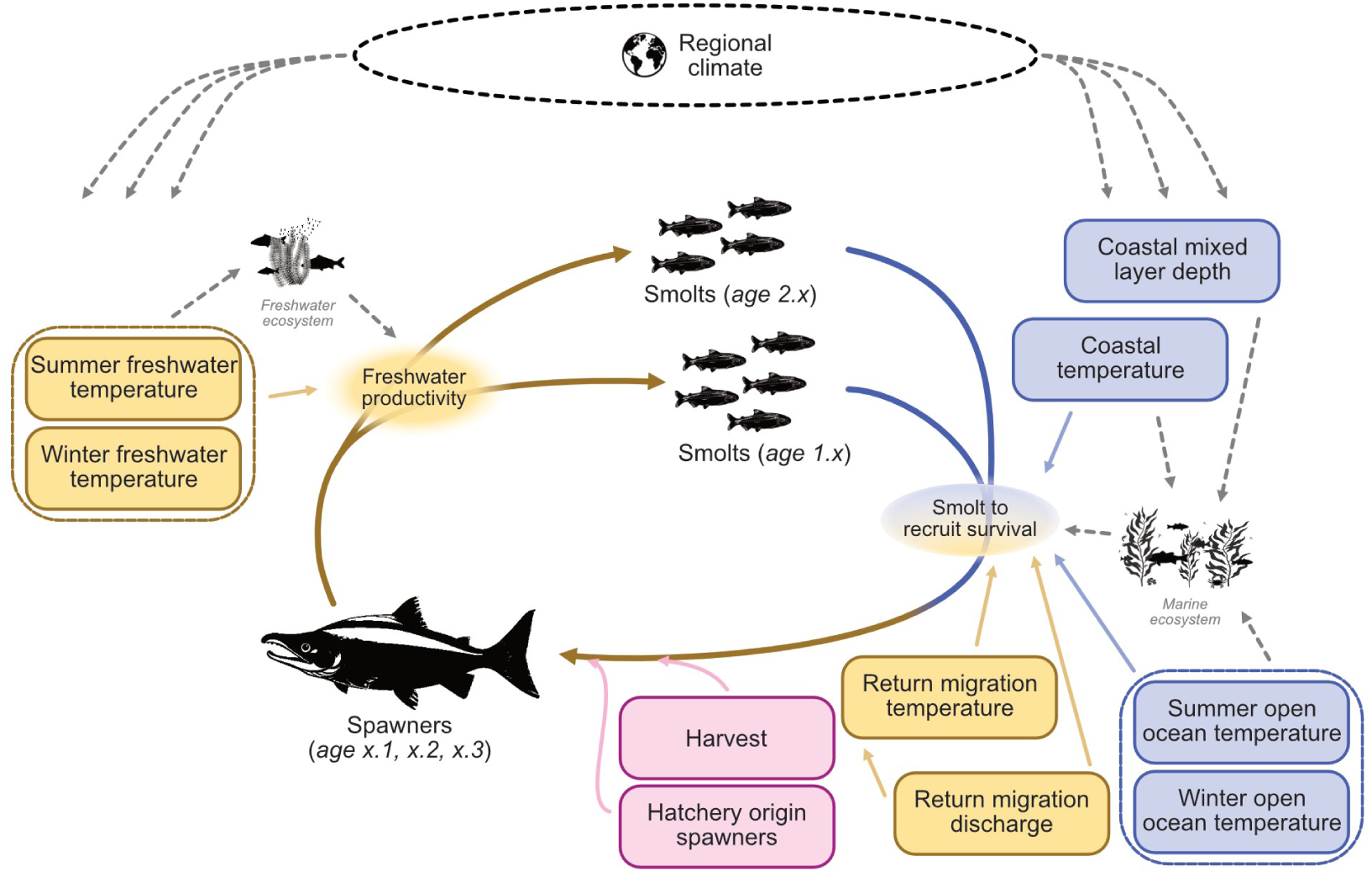
Conceptual model representing the sockeye salmon lifecycle and hypothesized environmental covariates associated with survival at freshwater and marine life stages. Arrows represent assumed causal pathways included in the analysis. Yellow shading denotes freshwater covariates and life stages, blue shading denotes marine covariates and life stages, purple denotes changes to spawner abundance via hatchery supplementation and harvest, unshaded nodes represent unobserved variables. A full directed acyclic graph (DAG) representing the explicit causal assumptions is shown in Figure S2

### 2.3 Freshwater Variables

Freshwater data for all populations except the two Columbia River populations were obtained from the Pacific Climate Impacts Consortium (Schnorbus 2024). We extracted gridded stream temperature and discharge data, and lake temperature data, based on rearing lakes and river migration routes (Figure 1). Historic data were observation-driven gridded data (gridded model data corrected by observation data) from 1981 to 2012. After 2012 data were missing and imputed in the model based on corresponding PCIC Global Climate Model (GCM) data for the Representative Concentration Path-way 4.5 (RCP4.5) scenario to reflect the mean warming trend and variance (Figures S5, S6, S7, S8). For the Columbia River populations (Osoyoos and Wenatchee), river temperature data were derived from USGS Station 12447200 at Malott, WA. Missing data were imputed using river temperature data and a model correlating air and freshwater temperature (Stiff and Thompson 2026). River discharge data for Osoyoos and Wenatchee lakes were taken from EnvCan Station 08NM127 and USGS Station 12462500. Lake surface temperatures were taken from EnvCan Station 08NM085 and missing data were derived from an air-to-water model (Stiff and Thompson 2026) for Oliver, BC. We calculated winter (October to March) and summer (April to September) indices to account for distinct effects of freshwater temperature on egg incubation and juvenile growth or survival, respectively. For the migration of adults returning to freshwater spawning grounds, we used freshwater temperature and discharge along population-specific freshwater migration routes between June to August (Figures 1, S4). The uncertainty of imputed data is shown in the supplement (Figures S5, S6, S7, S8). Due to large differences in discharge and temperature among rivers, river migration data were standardized per population, but lake rearing data were standardized globally. To reflect the assumed causal dependence of river temperature on discharge during upstream migration (Figure S2), return migration freshwater temperature (*FT* ^um^) and discharge (*FD*^um^) in a given year (*y*) were linked within the joint likelihood lifecycle model using relationships with population (*p*) specific intercepts (*α*) and slopes (*β*):

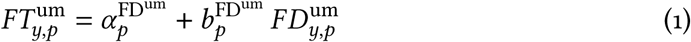

### 2.4 Marine Variables

We used model data for multiple sockeye salmon marine habitats, which span the continental shelf to offshore waters of the North Pacific Ocean. The Hindcast of the Salish Sea model (HOTSSea, resolution 1.5 km; Oldford et al. 2025) provided gridded ocean temperature data for the Salish Sea, where Fraser River sockeye salmon enter the marine environment. The British Columbia Continental Margin model (BCCM, resolution 3 km; Peña et al. 2019) is lower in grid resolution than the HOTSSea model, but covers a larger geographic extent along the west coast from the Columbia River to the Skeena River in northern BC. The Copernicus Global Oceanic and Sea ice Reanalysis model (GLORYS12, resolution 8 km; Lellouche et al. 2021) was used for the open ocean region and the northern coastal regions from the Skeena River mouth to Kodiak Island, Alaska. Gridded marine data were extracted from coastal regions bounded by Kodiak Island, each populations ocean entry location, and the edge of the continental shelf (i.e., 500 meters depth; Figure 1). The open ocean region boundaries were 170°W, 145°W, 60°N and 45°N, and a depth of greater than 1000 meters to exclude the continental slope. Temperature at five meters depth and mixed layer depth (MLD) were included as marine indicators in our analysis, capturing variability in thermal conditions and upper-ocean structure that influence habitat quality, productivity, and salmon survival. Temperature and MLD data were extracted for the coastal migration route from April to June in the year of ocean entry (Figure S4). For the open ocean rearing phase, temperature was integrated from July to September for the summer and January to March for the winter each year, matching marine rearing times. All marine data were averaged over the respective spatial and temporal domains, coastal and open ocean data were standardized globally for all populations. Missing temperature data were imputed based on corresponding Sea Surface Temperature data from the Extended Reconstructed Sea Surface Temperature model (ERSSTv6, resolution 2°x2°; Huang et al. 2025) (Figure S1), missing regional MLD data were imputed from MLD data for proximate regions and the regional temperature to MLD relationship with uncertainties (Figures S9, S10, S11, S12).

### 2.5 Process Model

2.5.1 Spawners to juveniles

We assumed the process model defining the freshwater stage sockeye salmon population dynamics followed a Beverton-Holt (BH) stock-recruitment relationship. The BH formulation is widely used in salmon population models because its parameters have direct biological interpretations in terms of productivity and capacity, and because it readily accommodates stage-specific recruitment within a broader life-cycle framework (Moussalli and Hilborn 1986). Fry abundances *F* were modelled as a function of spawner abundances *S* using density-independent *α* and density-dependent *β* parameters:

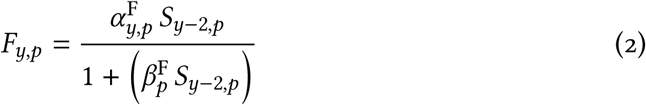

given each year *y* and population *p*. Here, fry abundance is specified at the end of the second winter following spawning, prior to the first fish in the cohort outmigrating as smolts. We modelled the density-independent fry productivity *α*^F^ (fry produced per spawner) on the log scale with hierarchical intercepts and environmental covariates:

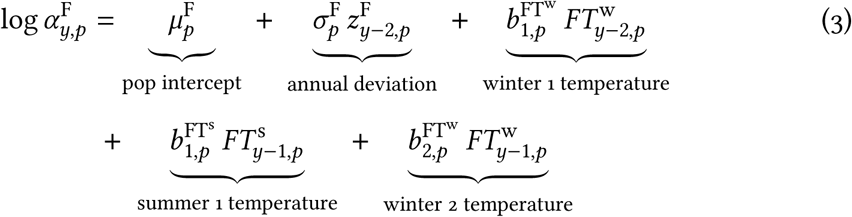

where 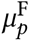 is the mean fry productivity for a population *p* with annual random-effect deviations 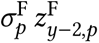. Mean fry productivity was modelled hierarchically across all populations and drawn from a global mean and standard deviation (Supplementary Table S3). Rearing temperature covariates were drawn from a hierarchy of global then domain-specific distributions as described in section 2.5.5. We included freshwater temperature effects from two winters *FT* ^w^ and one summer *FT* ^s^, with coefficients *b*^FTw^ and *b*^FTs^ , respectively. Parameters were indexed by the smolt year; the freshwater temperatures lag by smolt age to match the years of rearing experienced. After this period, the majority of fry migrate out to sea as age-1 smolts in the spring, while a smaller proportion migrate out to sea the following spring as age-2 smolts (details in following section). The density-dependent parameter 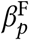 was modelled exponentially as a hierarchical parameter among populations (Section 2.5.5).

2.5.2 Freshwater holdover and age-2 survival

The relationship between fry and total smolt abundances was determined based on the population-specific proportion of fry that do not migrate as age-1 smolts (ℎ*old*), drawn from a hierarchical distribution. The proportion of fry *F* that migrate as age-1 smolts *Sm* was modelled as:

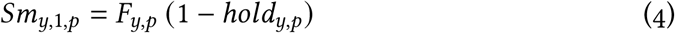

The number of fry that migrate as age-2 smolts is a function of ℎ*old* and a survival rate parameter *ε*^age2^ drawn from a hierarchical distribution:

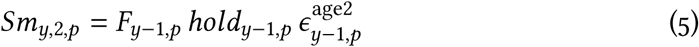

Survival of age-2 smolts 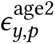 includes a prior year of freshwater temperature covariates for summer and winter before migration in the spring:

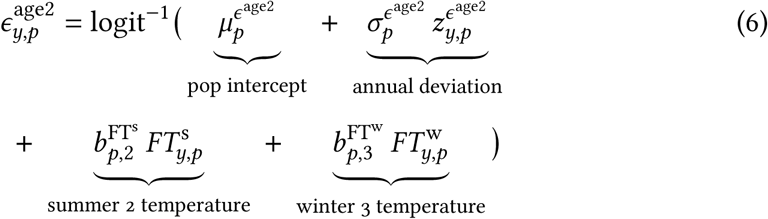

2.5.3 Smolts to returns

The effect of marine survival and return migration on smolts *Sm* to adult returns *R*, were modelled using a survival rate *α* specific to each year, population, freshwater age *f* and marine age *m*:

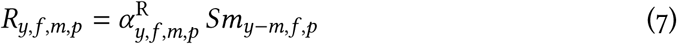

where freshwater and marine age class specific returns were linked back to a given smolt age class *f* equal to the current year less the return marine age*y*−*m*. The density-independent returns survival *α*^R^ is on the logit scale with hierarchical intercepts (Table S3) and environmental covariates:

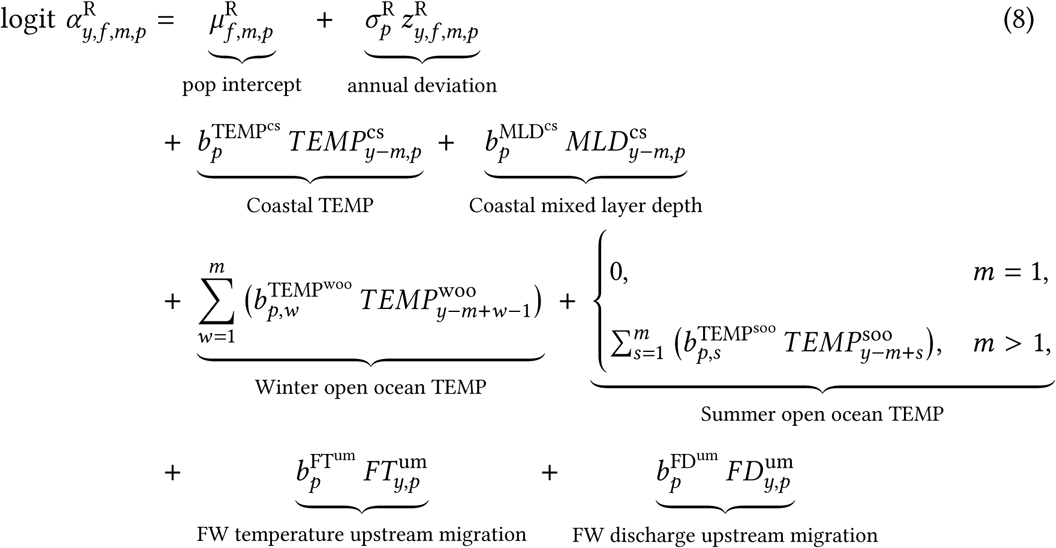

Where 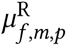 is the mean smolt to adult return survival for a given freshwater *f* and marine *m* age cohort from population *p* with annual deviations 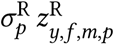. This mean survival parameter was modelled hierarchically across all populations for a given fresh- water and marine age cohort. Environmental covariates for the smolts to returns stage were included in the BH density-independent *α* parameter and drawn from a hierarchy of global mean and domain-specific means (Section 2.5.5 and Table S2). We included temperature (*T EMP*) and MLD along the coastal shelf (*T EMP*^cs^ and *MLD*^cs^, respectively) during the period from when salmon enter the ocean from the river mouth and migrate northwards along the coast to the Gulf of Alaska. We also included offshore temperature during the first ocean winter as a covariate (*T EMP*^woo^). For salmon that inhabit the marine environment for two or three years, we included additional summer TEMP (*T EMP*^soo^) and additional winter TEMP covariates. These are matched by rearing year specific 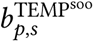 and 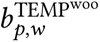 coefficients. Finally, we included freshwater temperature (*FT* ^um^) and discharge (*FD*^um^) as covariates during the upstream migration. Parameter indexing matched the return years, marine covariates were offset to match the corresponding year depending on marine age.

2.5.4 Returns to spawners

Total returns *R*^tot^, the sum of freshwater and marine age class specific returns per year, were deterministically related to the number of spawners *S* by accounting for fishing mortality *M*^F^ and hatchery returns *H*, where *R*^tot^ is the sum of salmon that return in a given year from different age cohorts:

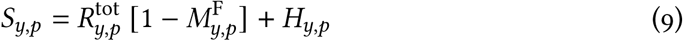

As noted above, we did not include data on mortality during freshwater migrations (i.e., en route mortality) because it was only available for two watersheds. Thus the effect of environmental covariates on recruit abundance integrates survival during marine and freshwater migration stages.

2.5.5 Hierarchical structure

Environmental covariate parameters were drawn from a global and domain specific hierarchy, priors are detailed in Tables S1, S2:

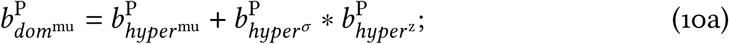

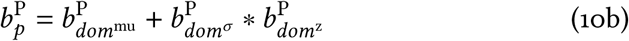

where 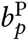 is the population specific parameter for covariate *P* and population *p*, 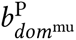 is the mean for the domain (*dom*) of a specific population with the associated *σ* and the population specific z, 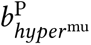 is the global parameter mean with its associated *σ* and domain specific z. All other parameters *P* had a simple hierarchy of global mean, sigma and population specific z (Table S3); the density dependent beta parameters *β* were drawn from an exponential distribution:

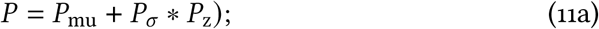

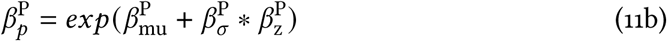

### 2.6 Observation Model

The lifecycle process model was fitted jointly with observation models for the different life stages in a state-space framework. Observed spawner data *S*^obs^ for each year and population were fitted to corresponding process model spawners *S*, using a negative binomial distribution with population specific overdispersion 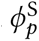:

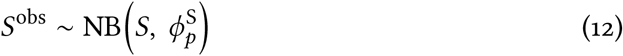

Juveniles were enumerated either as fry sampled in the lake (Osoyoos, Wenatchee, Great Central, Sproat, Chilliwack, Francois, Fraser, Shuswap, Quesnel and Babine Lake) or smolts sampled during the outmigration (Chilko, Tahltan and Tatsamenie Lake). Fry estimates in a given year *y* include all fish from brood year *y* − 1, which include those that will outmigrate as age-1 and age-2 smolts, as well as fry from brood year *y* − 2, which have held over and will outmigrate as age-2 smolts. Smolt data sampled in a given year *y* during outmigration include age-1 smolts from brood year *y* − 1 and age-2 smolts from brood year *y* − 2. Total smolts *Sm*^tot^ were the sum of age-1 and age-2 smolts in a given year. Fry and total smolt data were also fitted using a negative binomial distribution with population specific overdispersion:

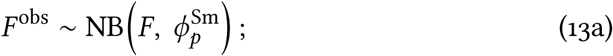

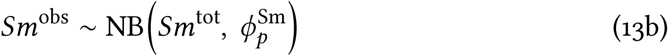

We fit both the observed smolt and return age-composition proportions *p^obs^* using multinomial distributions with logit-link functions:

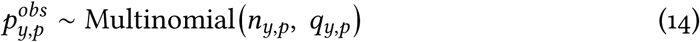

where *n* is the effective sample size that controls the contribution of each age-composition observation to the likelihood. Because sample sizes underlying the proportional age data were often unavailable, *n_y_*_,*p*_ was treated as an effective sample size representing the total information content and our relative confidence in each age composition observation, rather than the number of individuals sampled. The vector *q_y_*_,*p*_ = (*q_y_*_,1,*p*_, . . . , *q_y_*_,*A*,*p*_ ) represents the expected age composition for population *p* in year *y*, expressed as probabilities for each age class.

### 2.7 Model Performance

The combined process and observation model were implemented in Stan (Stan Devel- opment Team 2025) and cmdstanr (Gabry et al. 2025) in the R environment (R Core Team 2025). The model was fitted using an adapt delta of 0.99 and maximum tree depth of 11, over four chains (1000 warm-up and 2000 sampling iterations per chain, thinning of 2). Model convergence was assessed using trace plots, R-hat values and the bulk Effective Sample Size. To confirm that our model could reliably recover true environmental effects on productivity, we tested it on data generated by a simulation model in which the environmental effects were specified *a priori* as detailed in Supplementary Section 5.4.

### 2.8 Effect Size Calculation and Attribution of Environmental Effects

To facilitate comparison among environmental covariates and among populations, population-specific environmental effects were summarized as standardized effect sizes. Effect sizes were calculated as the change in productivity (returns per spawner) associated with increasing a given covariate from -1.5 to +1.5 standard deviations around its population-specific mean, while holding all other model covariates at their population-specific median values. For the summer and winter, freshwater rearing and open ocean temperature parameters, effect sizes for the multiple years were combined into one effect size. To quantify the relative importance of environmental covariates, spawner abundance, and random effects in explaining interannual variation in productivity, we decomposed model outcomes using Shapley values (Shapley 1952, Supplementary Section 5.5).

### 2.9 Projected Climate Impacts on Productivity

Future projections of sockeye salmon populations were generated by combining posterior distributions of demographic and environmental response parameters from the fitted Bayesian model with externally derived projections of environmental covariates for the time period 2041-2070. This approach propagates uncertainty from both estimated biological processes and environmental change into the forward projections, allowing responses to reflect parameter uncertainty and interannual climate variability.

We used an anticipated intermediate emissions scenario (Sarofim et al. 2024)) (approximated by SSP2-4.5/RCP4.5) for this analysis, which equated to increases in mean marine and freshwater temperatures of about 1.5-2°C while staying within historic ranges. Projected environmental covariates were obtained from multiple sources selected to represent freshwater and marine conditions relevant to salmon life history (Figure S4). Freshwater projections for all populations except the two Columbia River populations were obtained from PCIC GCM data for the RCP4.5 scenario, and included population-specific projections of freshwater temperature and discharge data. For Osoyoos Lake and Wenatchee Lake projection surface temperature data were derived from an air-to-water temperature model (Stiff and Thompson 2026) and projected air temperatures (Abatzoglou and Brown 2012), discharge data was fixed at the hindcast median because Columbia River flows are dependent on dam management decisions and cannot be projected based on GCMs. Projections of coastal MLD and temperature, and open ocean temperature were obtained from BCCM projections downscaled from CanRCM4/CanESM2 models (Peña and Fine 2024), and a custom set of statistically downscaled climate projections produced for this project (Actea San Francisco, CA) from CMIP6 data (Eyring et al. 2016). A challenge with statistical downscaling is that, unlike dynamical models, they do not explicitly solve the underlying physical and biogeochemical equations. As a result, they require validation not only against historical observations but also against independent high resolution dynamical simulations to assess whether their statistical relationships remain stable under future conditions (Dixon et al. 2016). Because no such dynamically downscaled projections exist for most of the Pacific salmon range, this important source of uncertainty could not be quantified for the custom statistical projections used here. Projection datasets were processed to match the temporal and spatial resolutions used in the historical analysis (i.e., population, river, or domain specific summaries) and standardized with hindcast parameters.

### 2.10 Artificial Intelligence Statement and Software

We used Microsoft 365 Copilot (GPT-5, 2025) to support Stan model development, troubleshoot Stan and R analysis code and for identification of potential causes of convergence issues. All AI-generated code and suggestions were verified, tested, and reviewed for logic and accuracy before implementation by the authors. Graphics in Figure 2 were generated using Affinity Designer (Serif, United Kingdom).

## 3 Results

### 3.1 Model Performance and Modelled Abundance Dynamics

The model converged with well-mixed chains and no divergent transitions, over 99% of draws had an R-hat below 1.01 and bulk Effective Sample Size above 400 (Figure S13). Modelled posterior predictions of spawner, smolt and return abundances generally tracked interannual patterns in the observed data, although agreement varied among populations and years (Figures S14, S15, S16). Observed values often fell within estimated uncertainty intervals, with greater uncertainty in periods with missing observations. Results from the simulation-recovery analysis indicated good parameter recovery (Figure S17). Across all populations and environmental covariates, the true parameter value fell within the estimated 95% credible interval for 97 of 104 parameters.

### 3.2 Environmental Variation

Overall, temperature covariates exhibited a warming trend from 1981 to 2024 (Figure 3), freshwater rearing temperatures ranged from 0-6.3°C in the winter and 3.9-18.3°C in the summer. Observed ocean entry temperatures in the southern watersheds ranged from 8.9-11.9°C, 9.9-11.9°C in the Fraser River watershed, and 6.6-9.7°C in the northern watersheds (data not shown). Coastal temperatures integrated along the migration routes showed alternating multi-year cold and warm periods, and relatively little variation among watersheds. Similarly, coastal MLD exhibited alternating periods of deep and shallow mixing with shallow mixing over the five year period 2019-2020. Observed winter and summer open ocean temperatures ranged from 4-6.1°C and 10.8-13.5°C, respectively, with a warming trend over recent years. Observed freshwater temperatures during return migration varied strongly among watersheds, ranging from 14.8-22.9°C in southern watersheds, 13.4-18.5°C in the Fraser River watershed, and 8-14.8°C in the northern watersheds. Across watersheds freshwater discharge spanned a wide range, from 4.8 m^3^s^-1^ in the Okanagan River (Osoyoos population) to 3524 m^3^s^-1^ in the Fraser River. Projections under the SSP2-4.5/RCP4.5 scenario for 2041-2070 indicate continued warming across most temperature covariates. Coastal MLD is projected to be relatively shallow in the future and river discharge during return migration is projected to decline for all watersheds.

**Figure 3:**
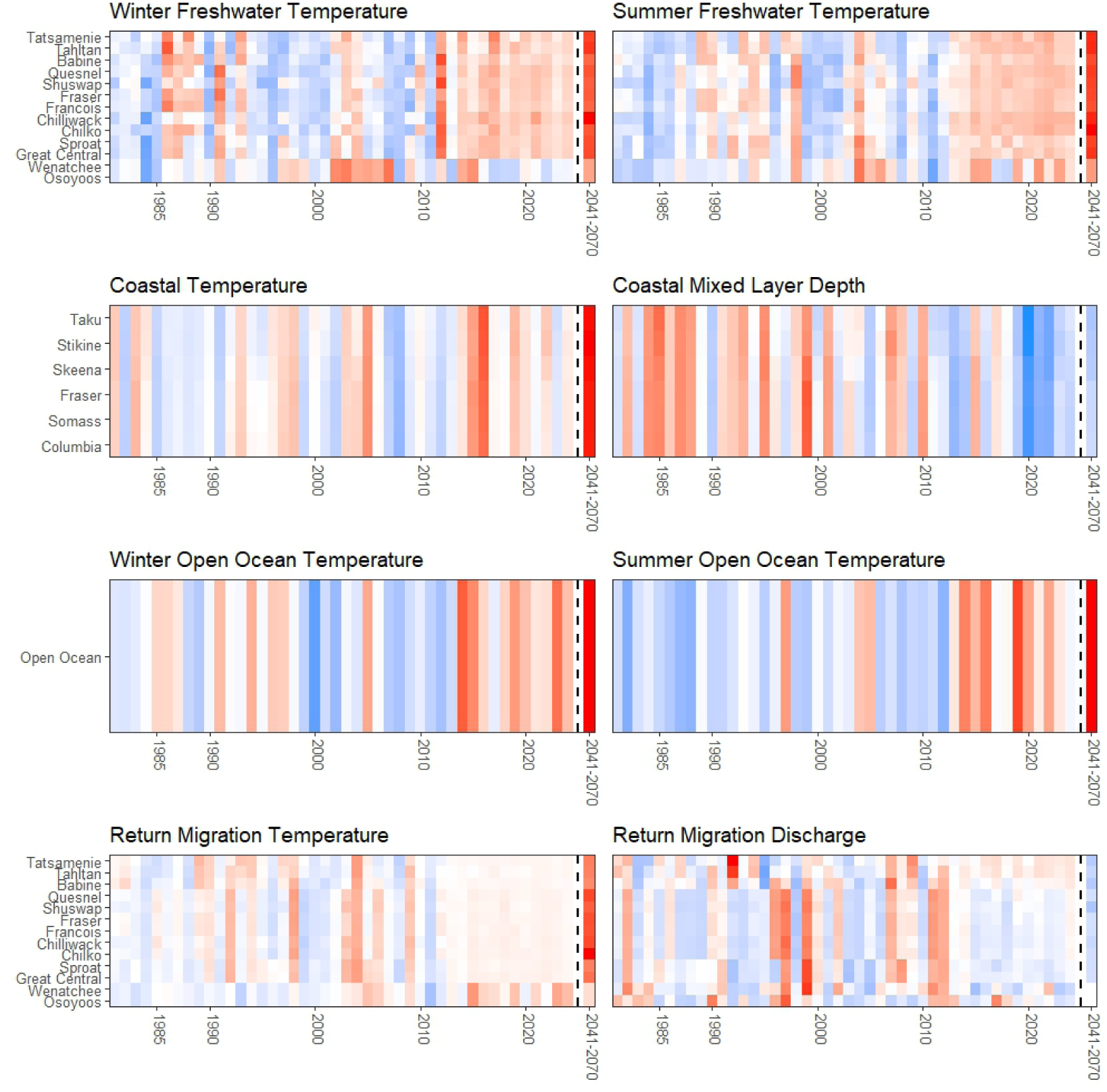
Standardized environmental covariates with missing data imputed, red indicates relatively high and blue relatively low values. Projection data for 2041-2070 are shown as means.

### 3.3 Environmental Response Parameters

The response parameters for winter freshwater rearing temperatures on productivity were centred on zero for nearly all populations (Figure 4, top row). In contrast, the parameters for summer freshwater rearing temperatures were negative in all domains and almost all populations except Tatsamenie, though the posterior distributions overlapped zero. Coastal temperatures had generally neutral effects for the Fraser River and northern domains, but a strong negative effect for the southern domain. Substantial among-stock variability in the coastal temperature parameters resulted in divergent patterns for specific populations within a domain, individual northern and Fraser River populations had relatively strong positive or negative relationships. Similarly the parameters for coastal MLD were consistently negative for the southern domain, with a neutral effect in the northern and Fraser River domains. Yet, a subset of northern and Fraser populations showed relatively strong effects of MLD, both negative and positive. The effects of winter and summer temperatures during open ocean life history periods were generally weakly negative with high uncertainty (Figure 4, third row). However, individual populations in the Fraser River and southern domains showed strong negative and positive responses. Return migration temperature parameters were generally negative across populations, with the exception of Tahltan in the north, but all posterior distributions overlapped zero. Parameters for the return migration discharge were negative for several southern and Fraser River populations though the domain-level and global distributions had considerable overlap with zero.

**Figure 4:**
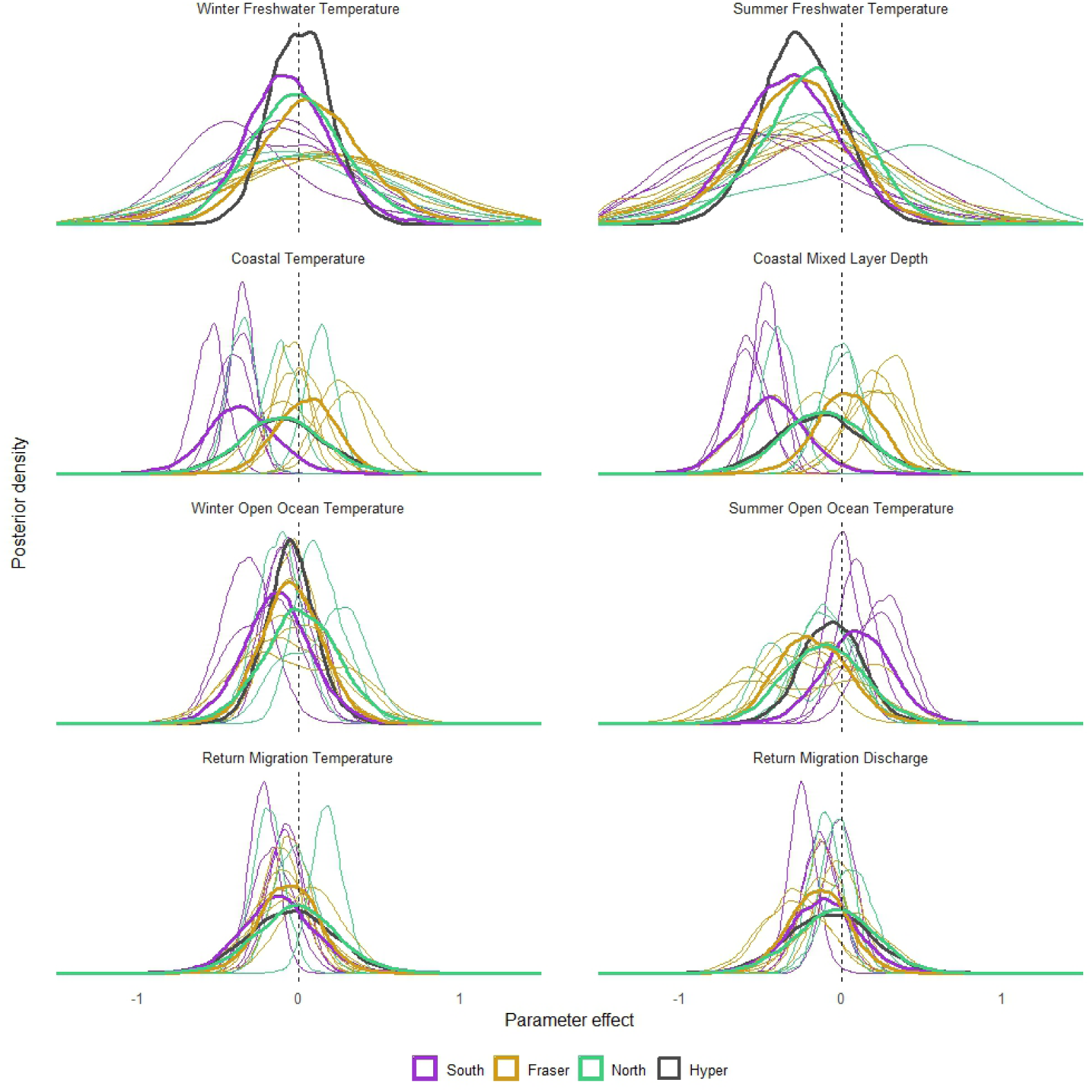
Distribution of parameters and across hierarchy levels, thin lines are populations-specific, thick lines are domain-specific means and black lines are global means. For summer and winter freshwater and open ocean temperatures, the distributions for multiple rearing years are combined. Colour coding by domains, Southern (purple), Fraser (gold) and Northern (Green).

### 3.4 Environmental Effect Sizes

Environmental effect sizes, expressed in changes in productivity, varied among populations and environmental covariates. Several of the predicted effect sizes were neutral (centred on or overlapping the 100% reference; Figure 5). Increasing freshwater rearing temperatures were generally associated with weak responses, although there was a tendency toward negative effects of increasing summer rearing temperatures (Figure 5, top row). These negative effects were most pronounced in southern populations, and in some cases, extended to winter rearing temperatures as well.

**Figure 5:**
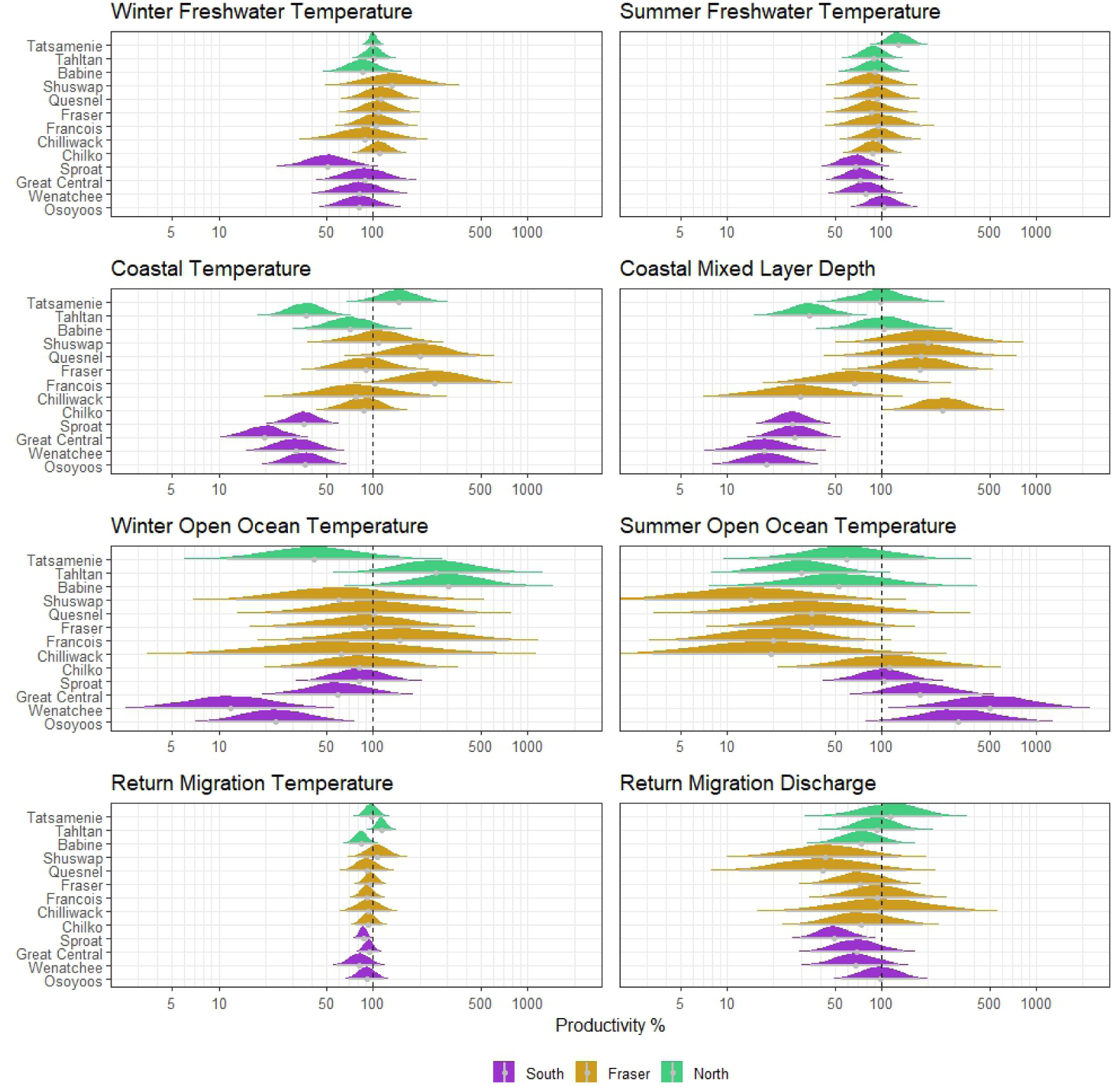
Change in productivity associated with a standardized effect of environmental covariates over a 3 SD variable increase. Distribution median and 95% interval are indicated in grey, 100% would represent no change in productivity. Colour coding by domains, Southern (purple), Fraser (gold) and Northern (green).

The effects of coastal temperature and mixed layer depth were much more pronounced, but varied strongly across populations and domains (Figure 5, second row). Increasing coastal temperatures and deeper mixing were associated with strong decreases in productivity for the four southern populations as well as Tahltan (northern domain), with posterior distributions only including negative responses. In contrast, the Fraser and remaining northern populations showed more heterogeneous responses. Some showed predominantly negative or positive responses to coastal temperature and MLD, while others were neutral, with posterior distributions that spanned both decreases and increases.

Effect sizes for open ocean conditions were much less certain compared to the other environmental covariates, with all populations exhibiting wide posterior distributions that frequently spanned both negative and positive effects (Figure 5, third row). In general, the southern populations showed negative responses to warming winter conditions and positive responses to summer temperatures. Fraser populations tended to have neutral responses to increasing winter open ocean temperatures and strong but uncertain negative responses to summer temperatures. Northern populations showed divergent responses to winter open ocean temperatures, with positive responses for Tahltan and Babine and negative for Tatsamenie. In contrast, all northern populations showed negative responses to increasing summer open ocean temperatures.

Responses to elevated return migration temperatures tended to be negative, but effect sizes were generally small compared to other environmental covariates (Figure 5, bottom row). High return migration discharge tended to have negative effects on productivity, although the posterior distributions included both negative and positive effects in all populations except Sproat.

### 3.5 Attribution of Interannual Variability

Shapley values indicated that, within the fitted model, combined environmental effects account for a substantial share of the variability in return abundance across populations (mean 31%; range 14-60%; Figure 6). In many populations, environmental contributions were comparable to or greater than those attributed to spawner abundance and random effects. Environmental contributions to return abundance were generally higher in southern and northern populations, and comparatively lower in Fraser River populations.

**Figure 6:**
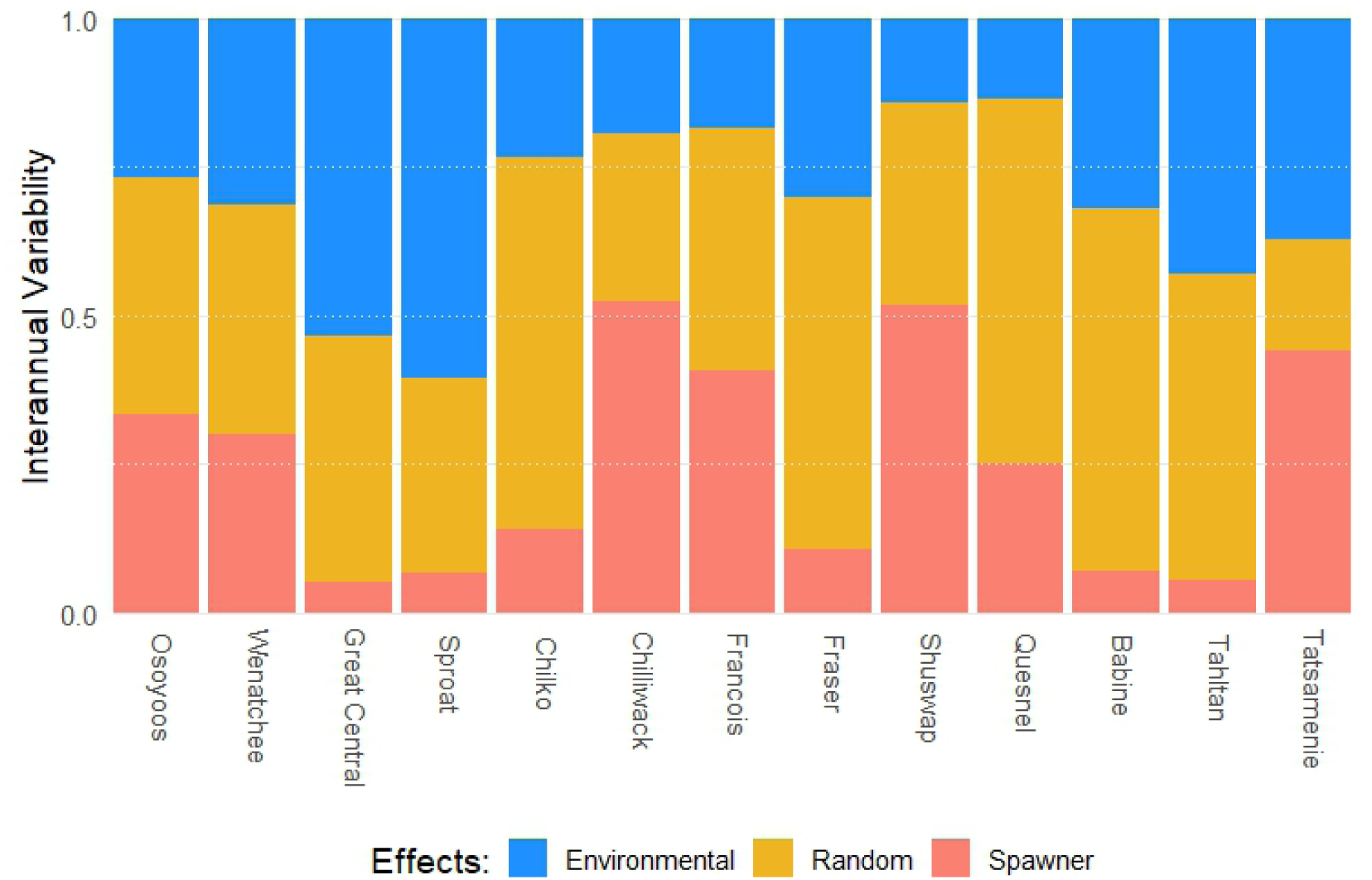
Relative contributions of environmental effects (blue), random effects (yellow) and spawner abundance effects (salmon) to interannual variability in return abundance for each population, based on Shapley values from the fitted model.

### 3.6 Future Projected Change

The combined projected changes in the environmental covariates included in our model were estimated to result in large but uncertain reductions in productivity between the historical baseline (1991-2020) and the future (2041-2070) SSP2-4.5/RCP4.5 scenario (Figure 7). Southern populations were projected to experience the largest declines in productivity, with posterior distributions entirely below the no-change reference. Projected productivity changes for Fraser River populations were varied, with negative central tendencies for Chilko, Chilliwack, and Fraser Lakes, and neutral responses centred near no-change for Francois, Shuswap and Quesnel Lakes. Projected productivity reductions were generally smaller for northern populations compared to those in other domains, and all northern populations had considerable posterior probability mass that included positive responses.

**Figure 7:**
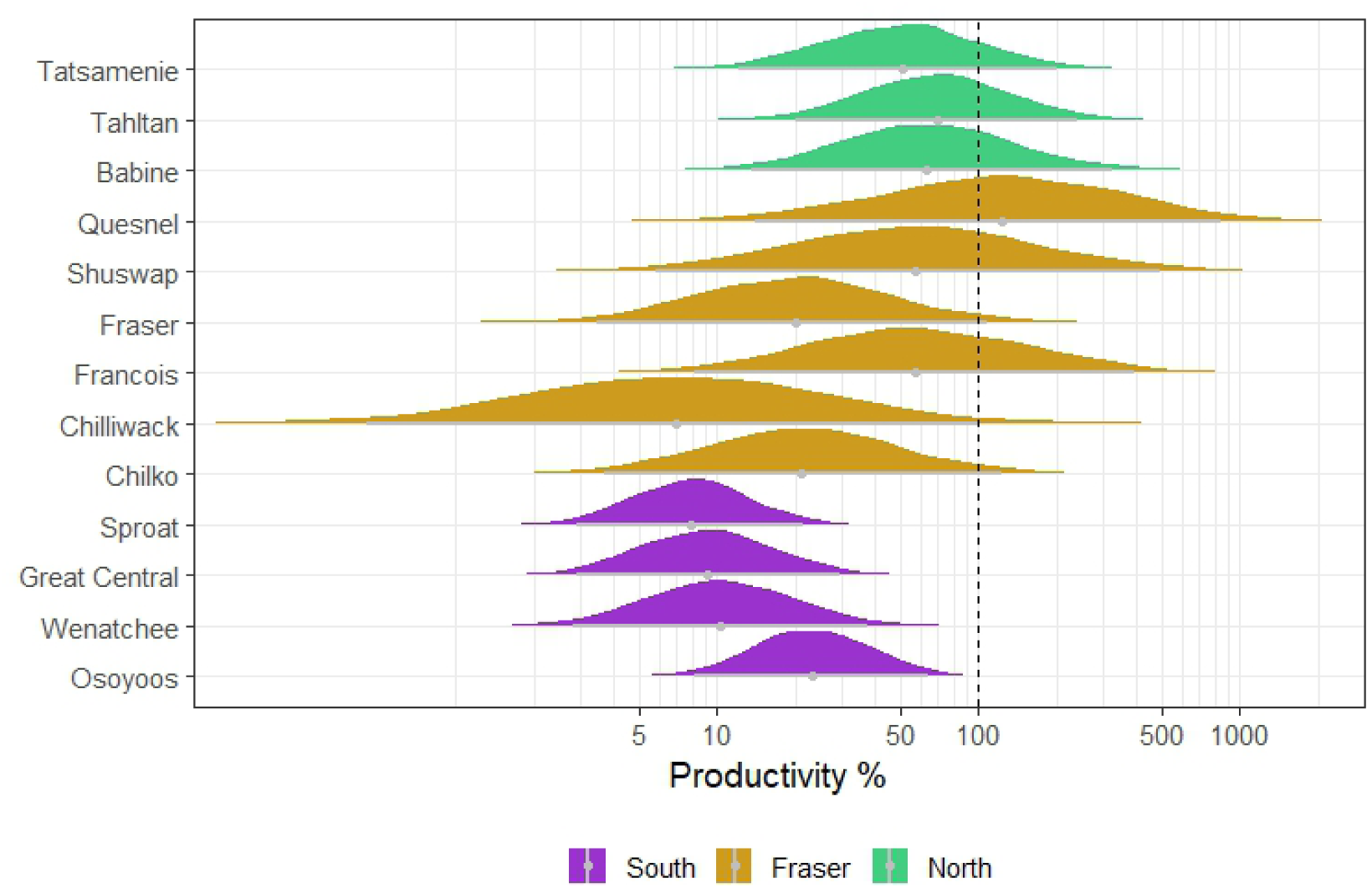
Combined effects of environmental covariates on productivity under the SSP2-4.5/RCP4.5 scenario for 2041-2070. Changes in productivity are scaled relative to the 1991-2020 reference period. Distributions represent the uncertainty of model parameters plus the interannual variability of covariates over the 30-year climatology, 100% would correspond to no change in productivity.

## 4 Discussion

Our analysis highlights that sockeye salmon productivity is sensitive to climate-driven change at different life stages across populations. Southern populations showed greater sensitivity to historical variation in coastal temperature, mixed layer depth (MLD), and freshwater discharge than Fraser and northern populations(Figures 4 and 5). Fraser populations showed strong sensitivity to open ocean summer temperatures. These patterns are reflected in the projected changes in productivity under future conditions, with southern populations expected to experience the greatest declines, Fraser populations showing mixed responses, and northern populations exhibiting modest declines. Together these findings demonstrate the value of explicitly modelling the full lifecycle of multiple populations and using a causal inference framework to assess life stage specific and combined environmental effects on salmon productivity.

Freshwater productivity, i.e., the number of juveniles produced per adult spawner, appears to be more sensitive to summer temperatures in nursery lakes than winter temperatures. For all domains and and populations, other than Tatsamenie Lake, our model estimated a negative effect of summer temperatures. This effect is strongest in three of the four southern populations, which also show negative responses to increased winter rearing temperatures. Temperatures in nursery lakes for southern populations often exceeded a commonly cited threshold of 15°C (Peacock et al. 2026) during summer and thus may be more vulnerable to further temperature increases in the future. While diel migration patterns provide salmon with cool, deep-water refugia from increased temperatures (Martins et al. 2012b), warming lake temperatures and associated hypoxic conditions lead to heat stress in juvenile sockeye salmon (Akbarzadeh et al. 2021), and the subsequent effects on their metabolism can affect smolt growth and survival. Furthermore, increased predation on juvenile salmon has been associated with warmer temperatures, potentially due to greater metabolic requirements of predators or changes in habitat use (Martins et al. 2012a).

Marine productivity is correlated with both local- and basin-scale indices representing conditions experienced shortly after ocean entry (Hobday and Boehlert 2001, Mueter et al. 2002, Kilduff et al. 2015, Wells et al. 2016, Connors et al. 2020). Our model focused on near surface temperature and MLD as two proxies for ecological conditions that may influence growth rates or predation-risk due to changes in the distribution of salmon and their predators. Both variables were integrated over large geographic domains and multiple months of marine residency to represent conditions during migrations along the continental shelf. We found that the survival of southern populations, as well as a subset of Fraser and northern populations, declined as waters warmed and MLD increased. Generally, increased coastal temperatures are thought to reduce salmon growth and survival via changes in prey availability, prey quality (Peterson and Schwing 2003, Ratnarajah et al. 2023), and metabolic rates (Thalmann et al. 2025), though they may also influence predator distributions (Emmett et al. 2006, Wells et al. 2016). A deep MLD reflects a well mixed upper water column, which affects phytoplankton and zooplankton productivity (Collins et al. 2009, Wolfe et al. 2016, Xue et al. 2022a, Suchy et al. 2025), and thus prey availability for juvenile salmon. For the southern populations migrating through the California Current, a deep MLD coincides with reduced upwelling driven nutrient transport (Xue et al. 2022b, Miller et al. 2026), further reducing productivity. The population-specific patterns we identified are generally consistent with evidence of latitudinal gradients in the effects of marine environmental covariates (Connors et al. 2020), which may reflect regional differences in salmon migratory behaviour, distinct physical oceanographic processes, or unique predator and prey communities.

Although recruitment variability is often thought to be driven by early marine survival, there is evidence that recent productivity declines in some populations are associated with increased mortality after the ocean entry year (DeFilippo et al. 2026). Open ocean winter and summer temperatures generally had uncertain effects on survival, though effects were stronger for a subset of populations. Specifically increased winter temperatures had a negative effect on several southern populations and a positive effect on several northern populations. In contrast, increased summer temperatures had negative, but uncertain effects on northern and Fraser populations, but generally positive effects on southern populations. The difference in effects among domains could be explained by population- or domain-specific marine distributions in the North Pacific. Although population-specific estimates of offshore distributions are not readily available, historical data have been used to highlight divergent patterns among species (Langan et al. 2024). Overall, open ocean temperature effects were more variable across populations than coastal temperatures, reflecting uncertainty in the spatial and temporal scale at which open ocean temperature effects on marine survival occur. Furthermore, uncertainty in the effect of offshore temperatures was likely increased by moderate collinearity between winter and summer temperatures in this stage, which limits the model’s ability to partition their effects. Rather than excluding one variable and attributing the effect to a single season, we retained both to reflect this uncertainty in a manner consistent with our causal framework. A final consideration is that competition (e.g., with Pink Salmon) represents an additional indirect effect of a warming ocean on sockeye survival (Connors et al. 2020, Ruggerone et al. 2023). As this pathway was not explicitly included in the model, it is implicitly included in the estimated total effect of open ocean temperature.

Increasing river temperatures and discharge rates during the return migration expose salmon to additional metabolic stress during an already demanding life stage (Eliason et al. 2011). However, the estimated relationship in our model indicates that higher discharge is associated with lower temperatures, such that their effects may partially offset one another. Temperature during upstream migration, especially in the Columbia River where temperatures often exceed a physiological threshold of 18°C (Peacock et al. 2026), can substantially decrease survival (Crozier et al. 2020). These effects may arise through increased metabolic demands (Eliason et al. 2011), as well as through elevated pathogen transmission and disease prevalence (Martins et al. 2011). River temperatures have a consistently negative effect for all populations, except the most northern ones in Tahltan and Tatsamenie Lake, but effect sizes are comparatively smaller than the impacts of temperature and MLD in the marine environment. This smaller effect size may be due to average conditions in most systems staying below commonly identified temperature thresholds (Crossin et al. 2008, Atlas et al. 2021), except for the Columbia and Somass systems. Our results are not consistent with the moderating effect of increased discharge rates on increased temperatures as recorded by Atlas et al. (2021). We see that some Fraser and southern populations show a substantial negative effect of increased discharge rates with peak discharge rates above 3000 m^3^s^-1^, which is supported by decreased survival of Fraser populations during the spring freshet due to increased energetic demands (Martins et al. 2011). The interacting effects of temperature and flow during return migrations are particularly likely to be location-specific and sensitive to features such as the presence of high- or low-flow barriers to migration.

We quantified the relative contribution of environmental covariates, spawners abundance, and other unexplained drivers of interannual variability (represented by random effects) on productivity using Shapley values. Generally the environmental covariates we considered accounted for a substantial portion of the model-explained variation in return abundance across populations. While spawner abundance contributes meaningfully, it does not dominate variability for many populations, consistent with evidence that recruitment is often not strongly associated with parental abundance or biomass across fish species (Szuwalski et al. 2015). In several systems environmental covariates explain a larger share of variation in return abundance than spawner abundance. In contrast, some Fraser River populations, including all of those with strong evidence of cyclical dynamics, show a greater contribution from spawner abundance relative to environmental covariates. Future assessments with population specific models incorporating environmental covariates can improve on solely abundance based spawner recruits models. Additionally, random effects also contribute substantially to the variability in return abundance in most populations, highlighting the presence of drivers of interannual variability that are not captured by the covariates in our model. These additional drivers (including predators, prey, competitors, surface wind and water turbulence, oxygen) could further improve model accuracy, but are data limited.

### 4.1 Projected responses to future change

The estimated responses to projected freshwater and marine temperatures, MLD, and river discharge show a distinct latitudinal gradient, with substantial declines in productivity for the southern and some Fraser populations, and relatively weaker responses in northern populations. The projected declines in productivity for southern populations are largely driven by warmer coastal temperature during the early marine phase and are further compounded by negative responses to warmer summer freshwater temperatures during the juvenile stage and to warmer winter temperatures during offshore residence. In contrast, a shallower MLD and warmer summer open ocean temperatures are projected to have favourable effects for southern populations under future climates, which partially moderate the projected decline. Additionally, freshwater temperatures in the Columbia and most Fraser systems are projected to regularly surpass high exposure temperature thresholds during river migration, further decreasing productivity.

The divergent projections for the Fraser populations reflect the relatively weak and variable estimated effects of environmental covariates in these systems. Because environmental covariates explain comparatively little interannual variation in populations such as Shuswap, Chilliwack, Fraser, and Quesnel Lake, projected responses are less pronounced and less consistent than in populations where these covariates were estimated to be stronger drivers of return abundance. This divergence may reflect lower sensitivity to environmental variation in these systems; however, it more likely suggests that our model does not include some relevant environmental covariates or that the common spatial or temporal scale of these covariates is not appropriate for all populations e.g., (McKinnell et al. 2014).

Northern populations generally showed the most favourable projected outcomes, their freshwater habitats and ocean entry areas are sufficiently cold that they are not expected to experience stressful conditions under a moderate warming scenario by 2041-2070, this matches the takeaway in previous projections (Stiff et al. 2018). Salmon populations at greatest risk from future environmental change are those that use habitats with large anticipated changes, show strong functional responses to environmental covariates, and occupy habitats in which stressful conditions compound throughout their lifecycle (Crozier et al. 2021).

The described projections are conservatively based on a moderate climate change scenario under emission mitigation (SSP2-4.5/RCP4.5). They should be interpreted as reflecting changes in productivity associated with the given environmental covariates under these conditions. Other drivers, including management interventions and additional environmental processes, may influence future outcomes and could lead to larger or smaller changes in productivity than those projected here. Yet, the findings hold lessons for future management decisions, there is a general latitudinal gradient in populations’ vulnerability to environmental covariates and interventions will be most effective if they target population specific life stages with disproportionate impacts on productivity.

### 4.2 Assumptions and Interpretation of Estimates

Our estimates depend on three critical assumptions: 1) that our directed acyclic graph (DAG) appropriately represents the causal pathways to each life stage; 2) that the environmental covariates used in the model capture historical variation in conditions experienced by each population during the corresponding life stage; 3) that the effects of historical and projected environmental change on productivity can be reasonably approximated by stationary, linear relationships.

Our DAG is simplified and attempts to represent the essential processes that influence salmon productivity with available data. It provides a transparent framework for identifying variables that must be accounted for to estimate the effects of interest and clarifying the assumptions underlying parameter estimates. By making these assumptions explicit, the DAG allows readers to assess how inferences might differ under alternative causal structures. A more detailed discussion of the DAG structure and associated assumptions is provided in Supplementary Section 5.1.

Clearly defining environmental covariates within our statistical model was particularly challenging because migrating fish experience conditions integrated over time and space. We addressed this challenge by averaging data over spatial and temporal domains based on published, data-driven life stage timing windows (Grant 2019, Lan- gan et al. 2024, Wilson and Peacock 2025) and expert driven decisions, as described in our Methods. However, averaging inevitably removes variation that may be important for determining salmon responses, and any mismatch between the spatiotemporal windows used in the model and the times or locations that most strongly regulate recruitment would dilute estimated effects. In addition, obtaining environmental covariates that span the spatial and temporal extent of our study required us to source several downscaled models and impute missing data, which introduce additional uncertainty that we were unable to propagate through our model.

We represented the effects of our environmental covariates on salmon productivity using stationary linear functional relationships. While the linear relationship simplifies the model, it likely does not accurately capture the true functional forms of some survival–environment relationships, particularly for temperature. The physiological effects of temperature on organisms are known to follow unimodal thermal performance curves, with performance declining beyond a thermal optima (Mayer et al. 2024, Arnoldi et al. 2025). We did not implement explicit nonlinear temperature responses here, in part because temperature effects are also operating indirectly through ecosystem processes that influence food availability and quality, and not only directly through salmon metabolism. The exception is in the upstream migration life stage, during which salmon have ceased feeding. For upstream migration, we explored threshold-based formulations to reflect declining survival at high temperatures. However, these formulations did not adequately capture expected responses, likely because migration temperatures are averaged over time and space. Similarly, the assumption of a stationary relationship simplified the model, and our data from outside a defined era of e.g. 1989-2012 (Litzow et al. 2018) was limited. Furthermore, assuming a nonstationary relationship would hinder our ability to make projections. These assumption have important implications for the projections, particularly when they require extrapolation beyond historically observed conditions. In populations where temperature effects are estimated to be positive, this reflects responses within historically observed temperature ranges and historical ecological relationships, rather than a direct beneficial effect of warming. For this reason, we interpret our projections as indicators of population sensitivity to future environmental change, rather than precise projections of future population status.

### 4.3 Conclusions

Our results highlight that climate-related drivers influence sockeye salmon productivity at both their freshwater and marine life stages. These responses are not uniform across populations or life stages and they can outweigh spawner-recruit relationships. This heterogeneity in responses underscores the importance of modelling multiple life stages, rather than relying on single-stage models and traditional spawner–recruit relationships. Echoing Champagnat et al. 2026, by integrating environmental effects on productivity within a causal lifecycle framework, our analysis provides a more mechanistic basis for understanding the impacts of a changing climate. Future changes in productivity will depend not only on the magnitude of environmental change, but also on how those changes interact with population and life stage specific sensitivities. Our results highlight the value of life cycle based assessments that integrate environmental and ecosystem covariates, and of management strategies that account for heterogeneity in population responses to environmental change.

## 5 Supplementary Material

### 5.1 Causal Inference Supplement

Estimating causal effects from observational data is challenging because environmental covariates often influence each other or are correlated because they share a common cause. The causal inference framework allows us to handle this complexity by making our assumptions explicit and linking those assumptions to the structure of our statistical model (Pearl et al. 2019). Central to this framework is a directed acyclic graph (DAG), which is a diagram that formally expresses our scientific understanding of how variables affect each other. The DAG then guides the construction of the statistical model; it identifies which variables must be included in the model (because we want to estimate their effects or because they are needed to remove non-causal associations), which variables to leave out (because including them would bias our estimated effects), and clarifies how to interpret the estimated effects. Because causal estimates depend on the assumed causal structure, formalizing those assumptions in a DAG makes them explicit and open to critique, allowing others to propose alternative structures and assess how changes might affect inference (Champagnat et al. 2026). This approach differs from the common practice of using multiple regression with model selection to identify predictive environmental covariates, which implicitly defines a causal structure where all covariates have direct effects on the response and are independent of one another. Such approaches are not designed for causal inference and can yield biased effect size estimates when the implicit causal structure is inconsistent with the hypotheses being tested (McElreath 2020).

**Figure S1:**
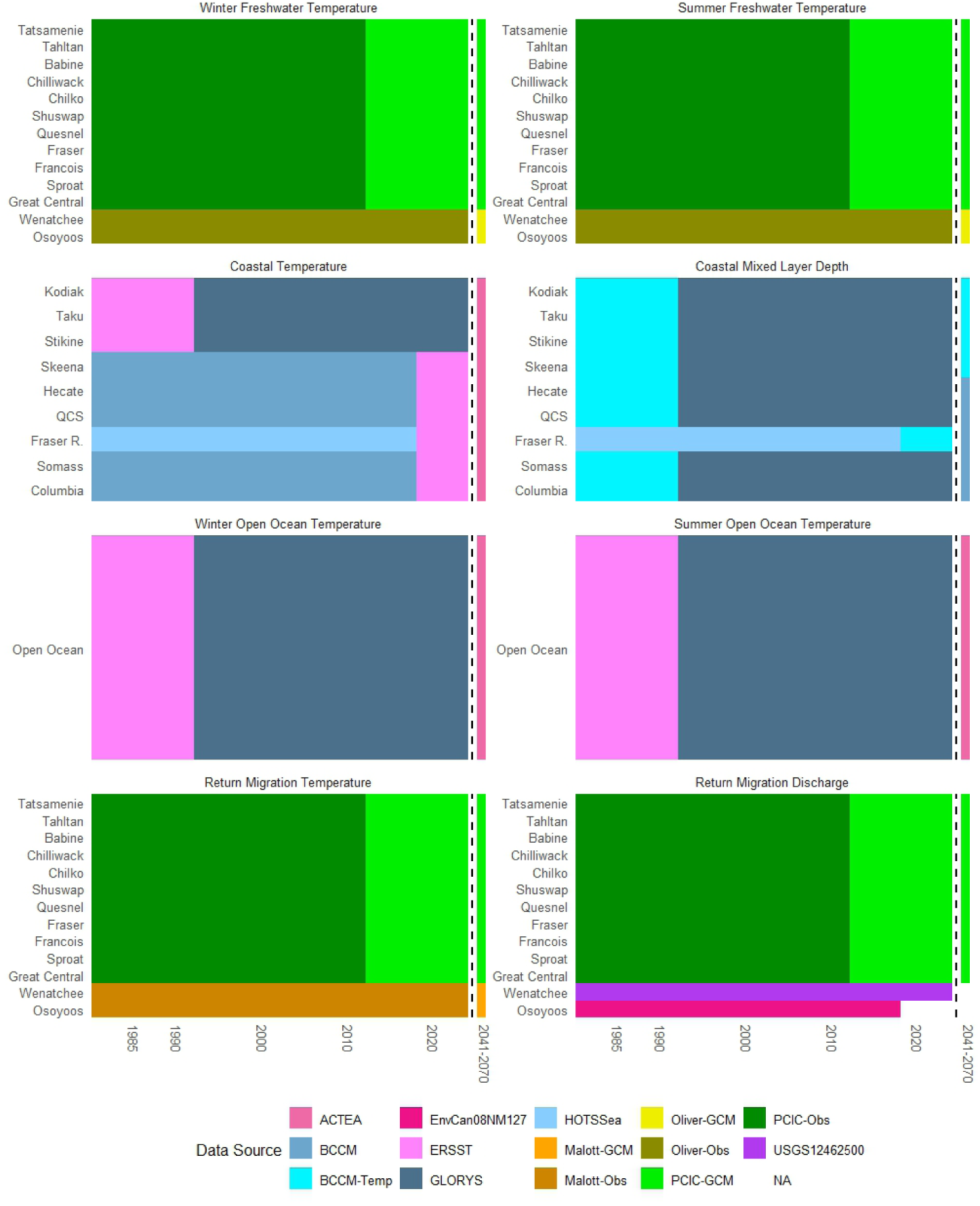
Data sources for the different data types by population or region. For missing data, ERSST data is used to impute missing near surface temperature data at 5 meters; BCCM-Temp refers to a combination of BCCM MLD and temperature data to impute missing GLORYS MLD data; for Malott, Oliver, and PCIC, data is primarily derived from observations-based data ”-Obs” and missing data are imputed from global change model ”-GCM” data. There were no projections for discharge data for the Columbia River Basin (NA).

**Figure S2:**
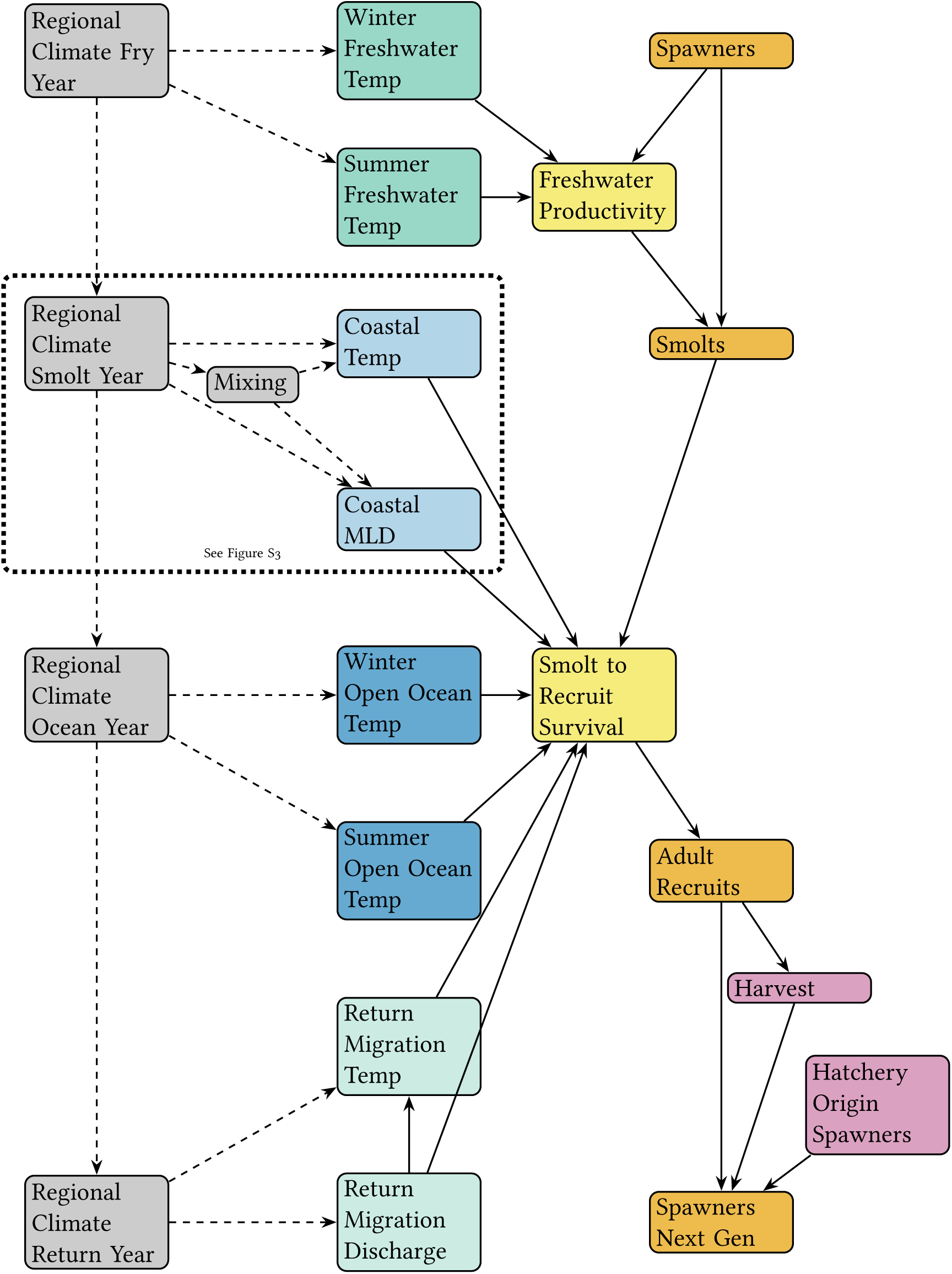
Directed acyclic graph detailing the model structure, variables and associated correlations. Grey boxes with dashed arrows indicate unobserved higher-level variables. Green and blue boxes are freshwater and marine covariates (MLD - Mixed Layer Depth, Temp-Temperature), yellow boxes are the process model covariates, orange and lilac boxes are the observation model covariates. The Coastal Temperature - Coastal MLD relationship (dotted frame) is detailed in Section 5.2 and Figure S3.

Figure S2 provides the detailed structure underlying our model. Because our goal was to estimate the impacts of climate-associated drivers, the DAG does not explicitly represent many known and hypothesized biological drivers of salmon ecology. Instead, it focuses on temperature, mixed layer depth, and river discharge and their direct and indirect effects on sockeye survival at different life stages.

Here, we assume that temperature in both the freshwater and marine stages influence salmon directly (e.g., via changes in metabolism), as well as indirectly, by altering the availability of prey. Because prey abundance lies downstream of temperature and is not included as a covariate, the estimated temperature coefficients should be interpreted as capturing the total effect of temperature, including both direct effects and indirect effects mediated through prey.

Many additional variables influence prey availability, such as nutrient availability, primary production, and competitors. If these variables are themselves influenced by temperature (i.e., mediators), as is often the case, then their effects are implicitly included in the estimated temperature effect. If they do not share causal pathways with temperature (i.e., if they affect prey availability or survival but do not influence temperature), omitting them from the model should not bias the estimated effect of temperature, although including them could increase the precision of the estimates by accounting for other causes of variation (Pearl et al. 2019). In contrast, variables that influence temperature and prey availability, or other drivers of survival, can introduce spurious correlations and bias our estimated effects if not accounted for.

For example, temperatures in the coastal migration life stage and both seasons of the open ocean life stage are correlated since they are influenced by the regional climate of the North Pacific. This is addressed in our model by including temperature covariates at multiple locations in the lifecycle, which blocks the non-causal association created by their shared dependence on regional climate. However, these temperature covariates remain a simplified representation of the continuous environmental experience of the fish. Thus, the estimated coastal temperature effect may be partially capturing the effect of conditions outside the spatial and temporal windows defined for the coastal migration stage.

### 5.2 Temperature and Associated Covariates

The non-causal associations described for temperatures are further complicated when we also consider the effect of other covariates. In the coastal environment, MLD is a determining variable for productivity. MLD defines the depth of the surface mixed layer of the ocean in which density is nearly uniform, as determined by turbulent mixing processes in the upper ocean (De Boyer Montégut et al. 2004). Changes in MLD influence nutrient availability in surface waters by regulating the degree of vertical exchange with deeper, nutrient-replete waters that support primary production. On the other hand, increased mixing associated with deeper MLD can reduce the sunlight received by primary producers, such that the net effect of MLD on primary production is nonlinear and often conceptualized as an ’optimal stability window’ (Gargett 1997). Variation in primary production, in turn, influences the abundance of zooplankton and forage fish that serve as prey for sockeye during their marine life stage; but a deeper MLD can also reduce grazing efficiency of zooplankton (Xue et al. 2022a), thus reducing the prey availability for sockeye. Figure S3 provides a more detailed articulation of the assumed causal pathways for productivity in the coastal migration life stage.

**Figure S3:**
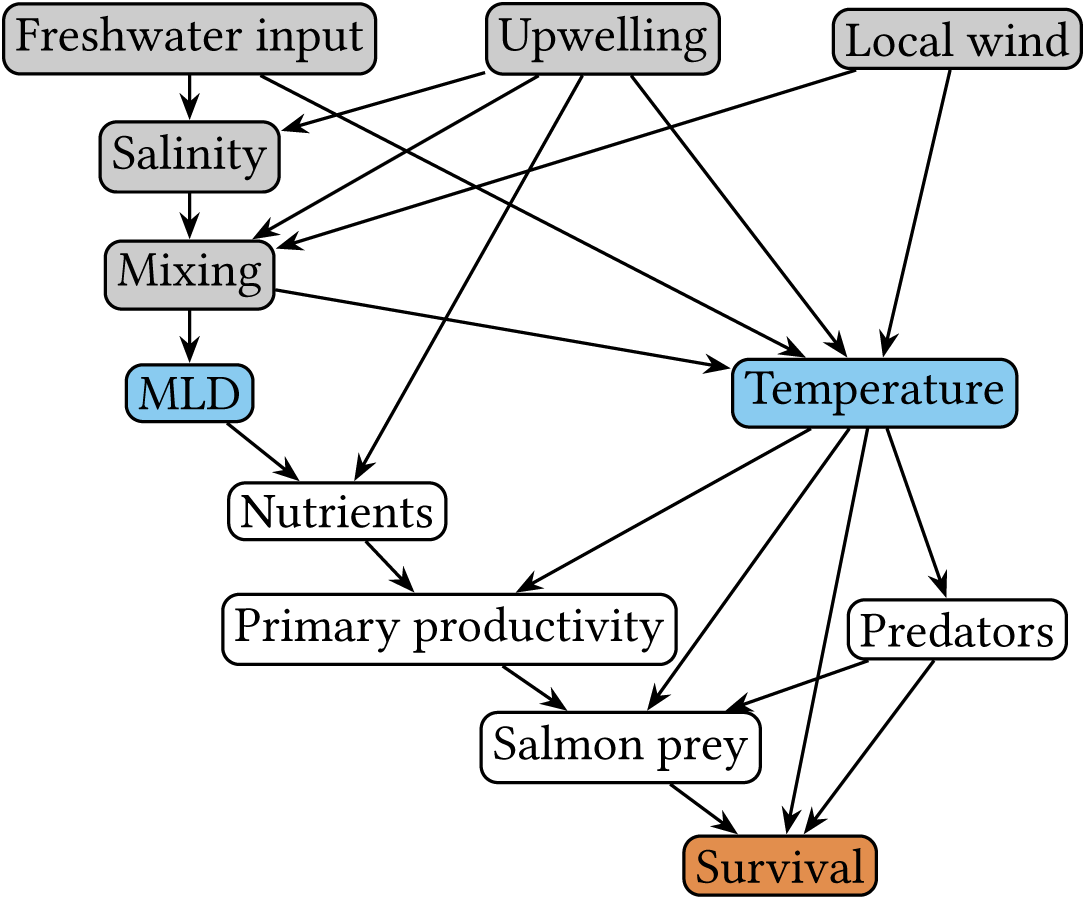
Directed acyclic graph of temperature and mixed layer depth effects on sockeye marine survival as well as a simplified set of associated causal variables. Orange: outcome variable; blue: covariates in our model; grey: upstream variables not included in our model; white: unobserved mediating variables.

Temperature affects metabolic rates (Kooijman 2000), therefore it is assumed to influence primary production, prey availability, predator abundance, as well as salmon survival directly. We do not assume that temperature is a primary driver of predator abundance, but we include this pathway to reflect the possibility that predators respond to temperature and to illustrate that conditioning on predator abundance is not required, or desirable, to obtain an unbiased estimate of the effect of temperature on salmon survival. Since we lack observations of nutrients, primary production, prey availability, and predator abundance, we cannot estimate their impacts on survival, or use them to distinguish the direct and indirect effects of temperature on survival. As a result, the estimated coastal temperature coefficient captures both its direct effect as well as all its indirect effects on the ecosystem. In contrast, there are no assumed direct effects of MLD on salmon survival and so its coefficient captures its indirect effects on the ecosystem (Figure S2).

Importantly, MLD and temperature are both influenced by upper-ocean mixing and its drivers (e.g., wind, upwelling, and freshwater input), which cause them to be correlated. Including both temperature and MLD as covariates of salmon survival blocks these shared-cause pathways and accounts for the correlation between them (Figure S3), but the drivers of mixing do not need to be included as additional covariates, provided they do not affect survival through other causal pathways. Coastal upwelling is the exception as it is assumed to influence nutrient availability directly, independent of its effects on mixing, MLD and temperature. This creates the possibility of associations between MLD or temperature and nutrients that are driven by upwelling rather than by those variables themselves, which could bias the estimated effects of temperature and MLD on survival. However, we expect that this bias should be relatively small since the effect of upwelling on available nutrients is further mediated by mixing, and this pathway is blocked in the model by the inclusion of MLD as a covariate. Including upwelling as an additional covariate would fully block this remaining pathway, but we were unable to identify an estimate at an appropriate spatial and temporal scale for this study.

For freshwater rearing environments we do not have estimates of mixing or stratification to match temperatures. Because of this, our estimates of temperature in these life stages are likely somewhat biased by the fact that we cannot control for correlations between temperature and salmon survival that are due to the types of shared processes outlined for the coastal environment. Nevertheless, the structure of the DAG and the variables at play are different in these ecosystems. In the rearing lakes in this study temperature is a primary characteristic of stratification (Stockner and Shortreed 1983) and the estimated effects of temperature should be assumed to also be including the non-temperature related aspects of mixing.

During upstream return migration, the assumed relationship between river discharge and temperature in the return migration requires consideration. Our model includes both discharge and temperature as covariates of survival. Including discharge ensures that we are estimating an unbiased effect of temperature. As a consequence, including temperature blocks the indirect effect of discharge on survival, such that the discharge effect parameter captures only the direct effect of discharge. However, because our joint likelihood model explicitly models the effect of discharge on temperature, we can use posterior simulations to estimate the total effect of discharge on survival, including both the direct and indirect pathways.

**Figure S4:**
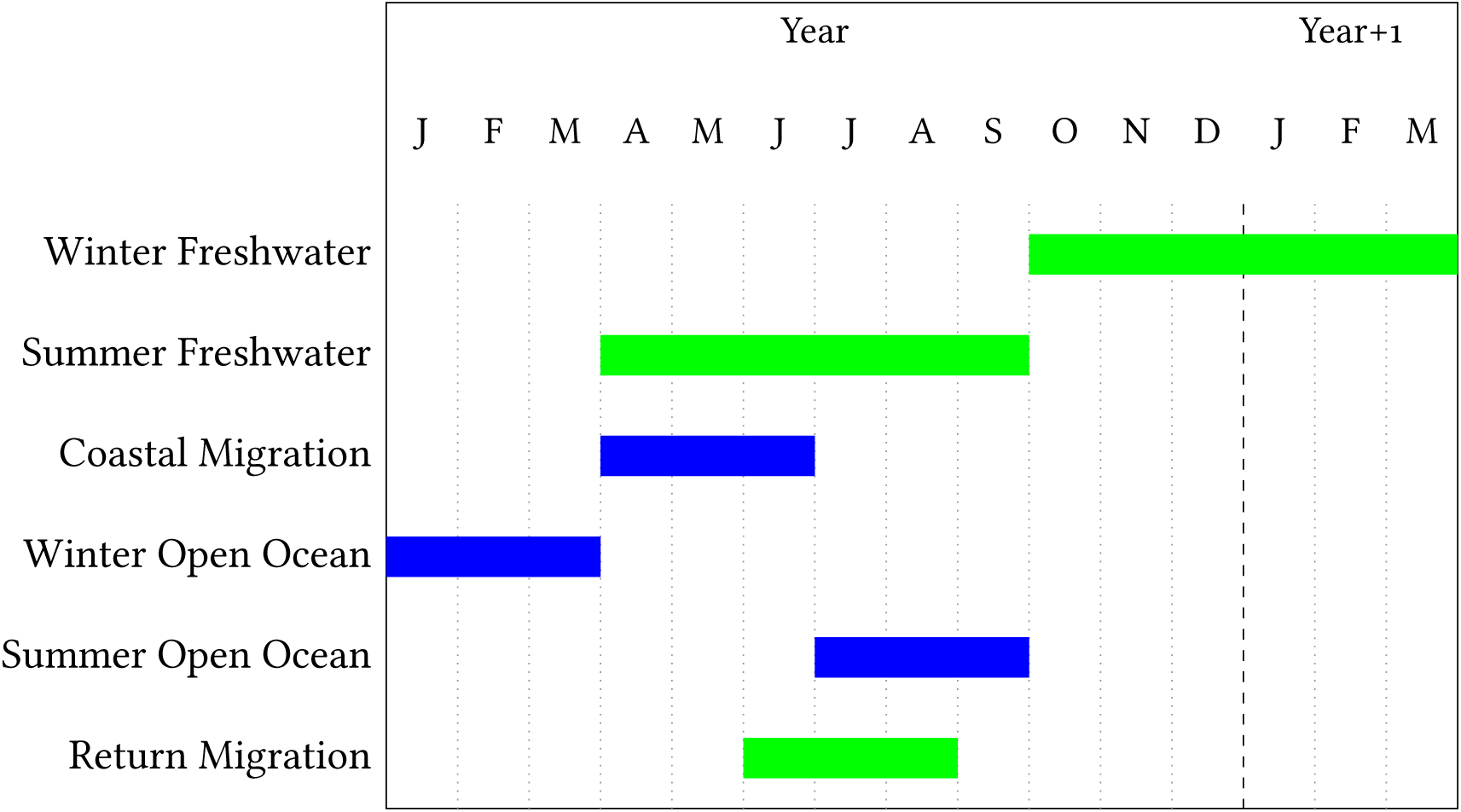
Chart showing the temporal windows used for environmental data integration across life history stages. Freshwater covariates are shown in green and marine covariates in blue. Letters indicate months of the year.

**Figure S5:**
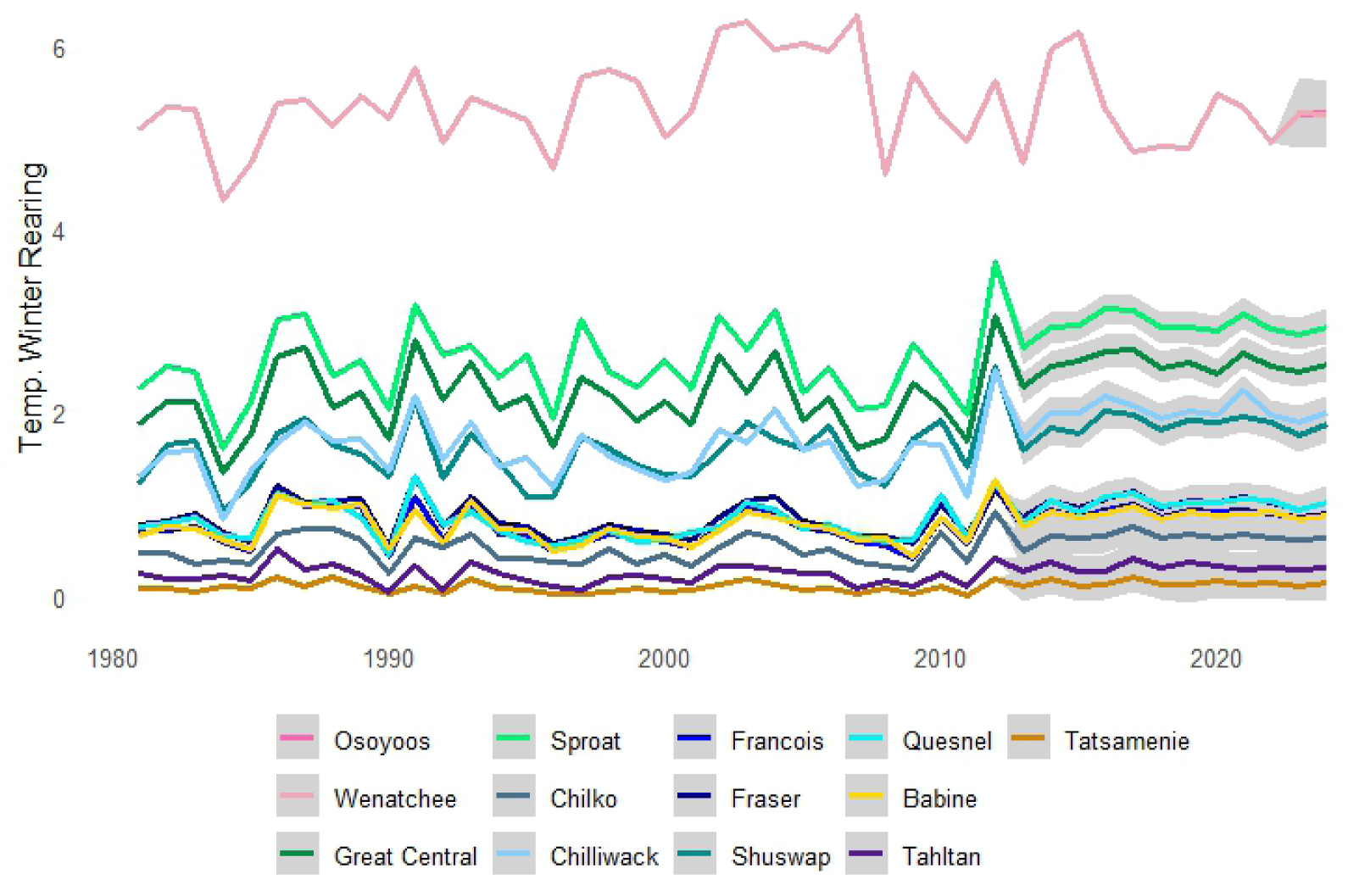
Winter Rearing Temperatures in °C, colours indicate the population. Imputed data shown with 50% credible intervals. Osoyoos and Wenatchee data are overlaid.

**Figure S6:**
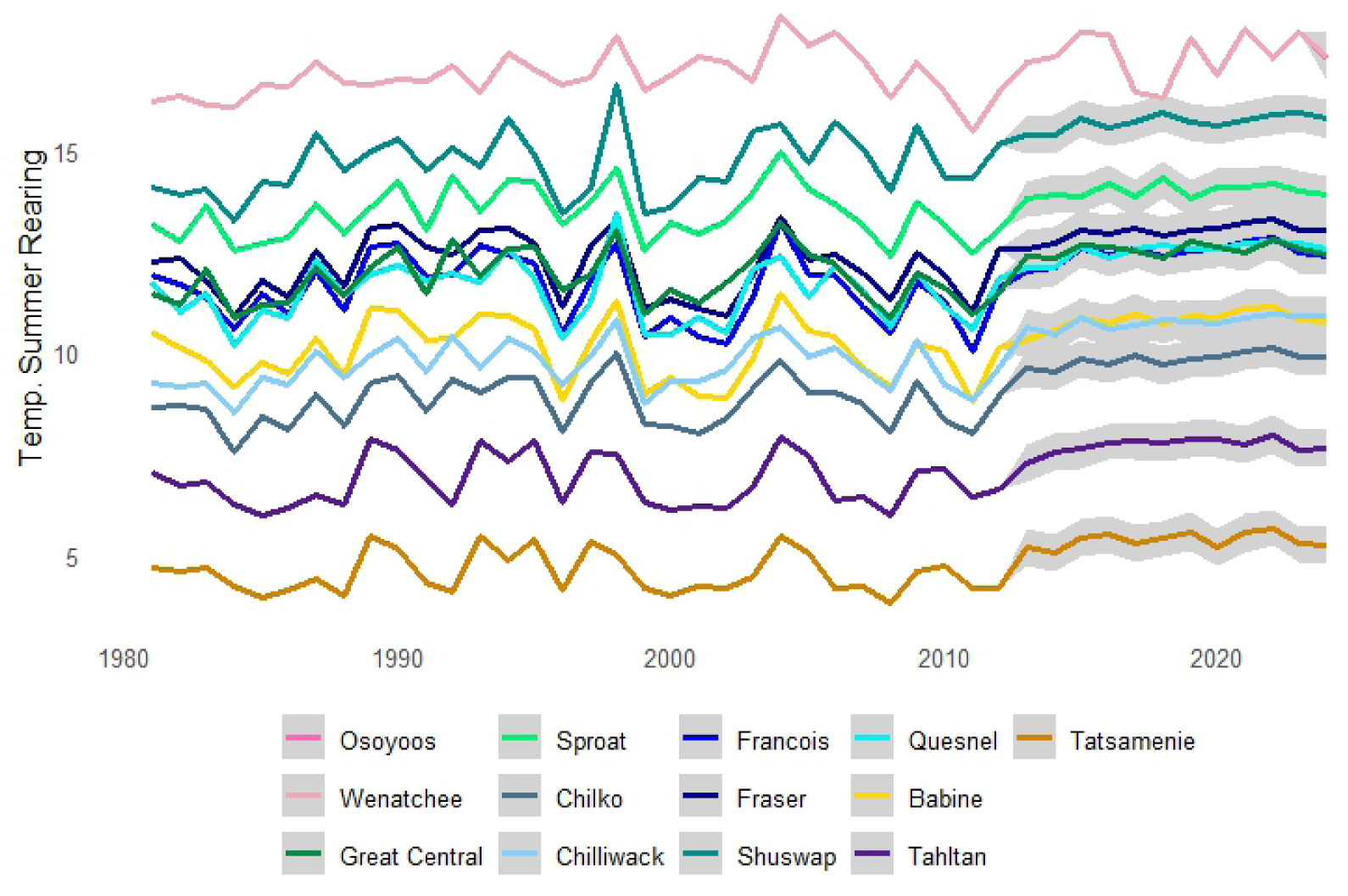
Summer Rearing Temperatures in °C, colours indicate the population. Imputed data shown with 50% credible intervals. Osoyoos and Wenatchee data are overlaid.

**Figure S7:**
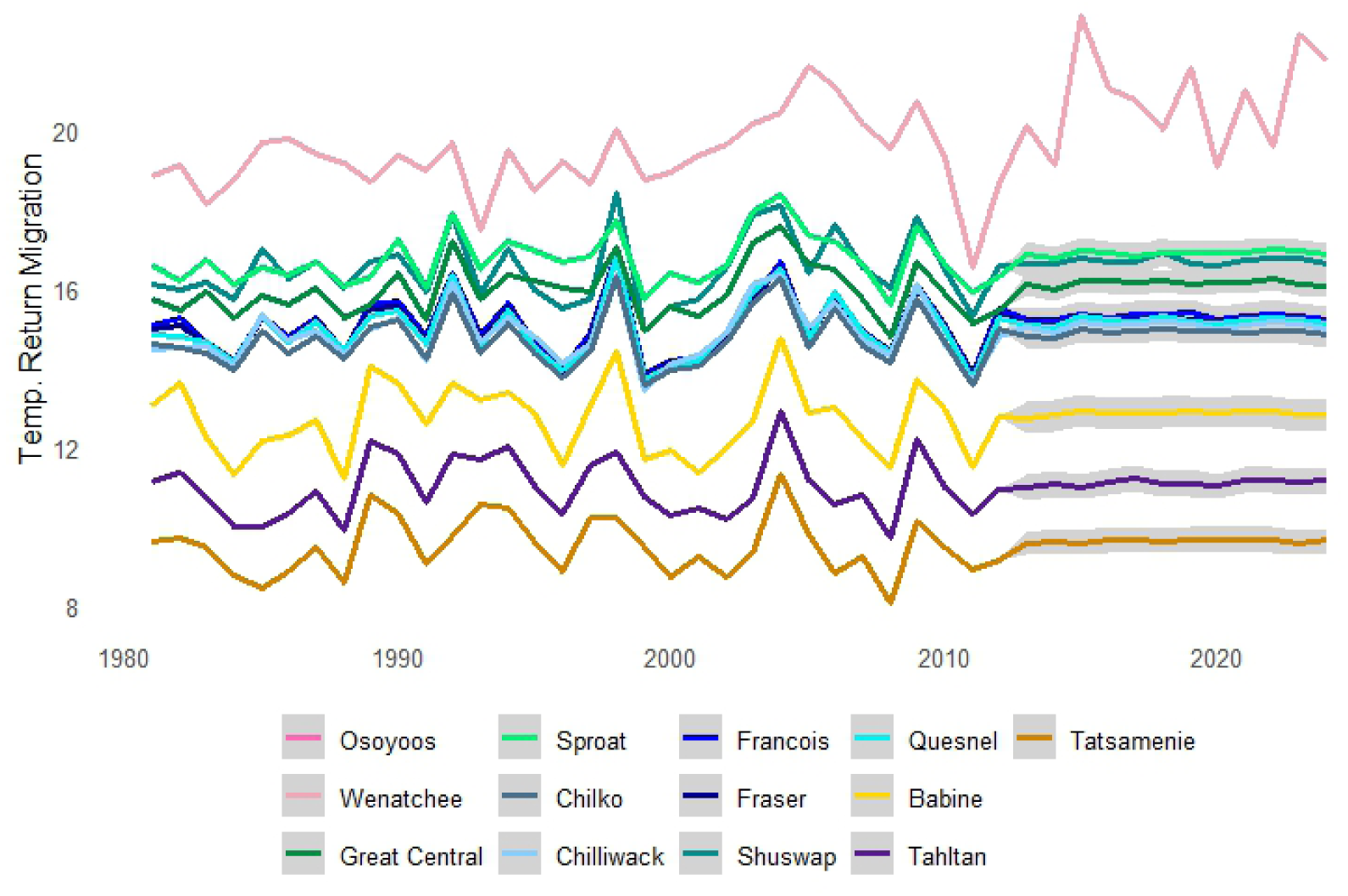
Return Migration Temperatures in °C, colours indicate the population. Imputed data shown with 50% credible intervals. Osoyoos and Wenatchee data are overlaid.

**Figure S8:**
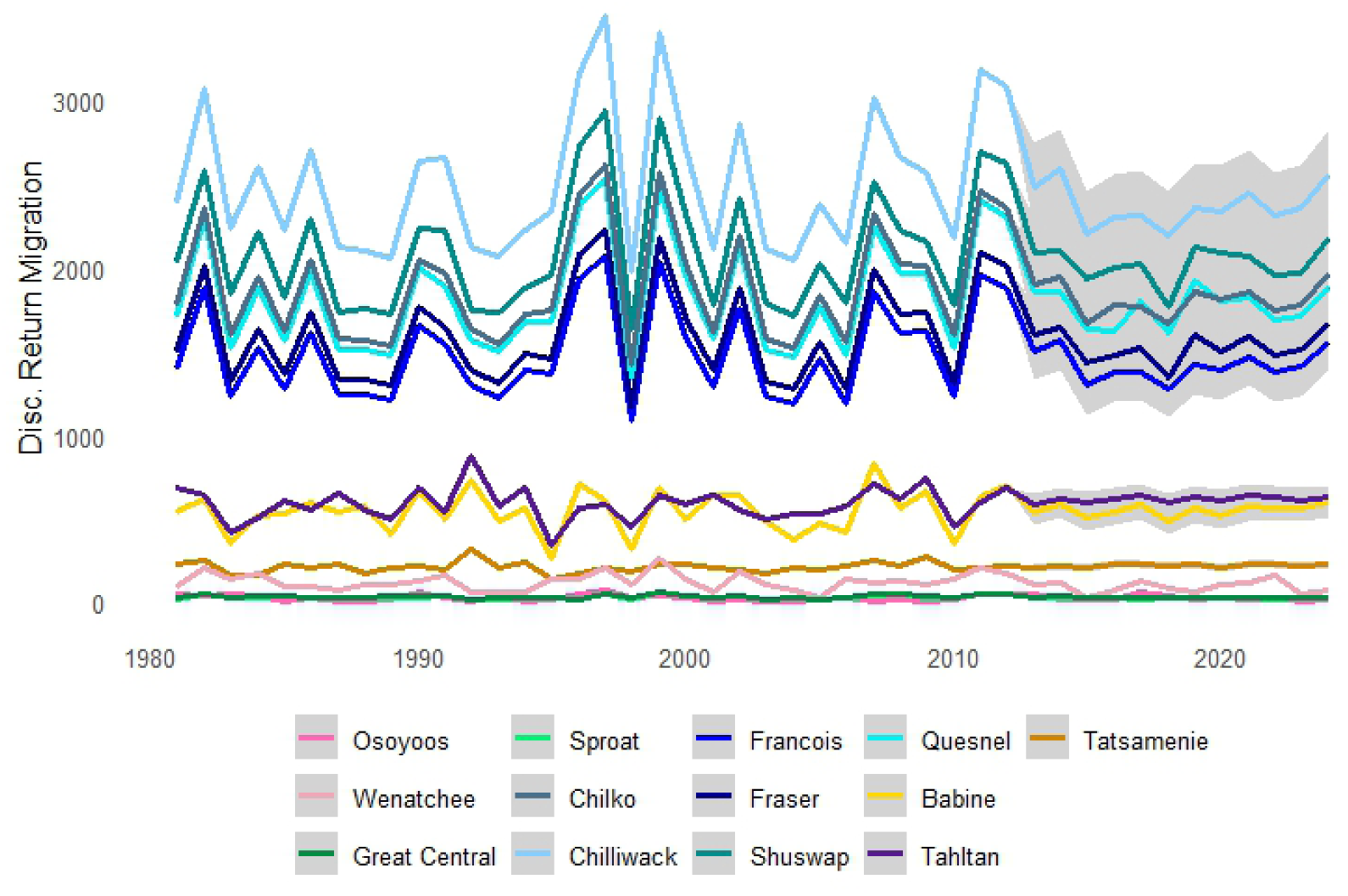
Return Migration Discharge rates in m^3^s^-1^, colours indicate the population. Imputed data shown with 50% credible intervals.

**Figure S9:**
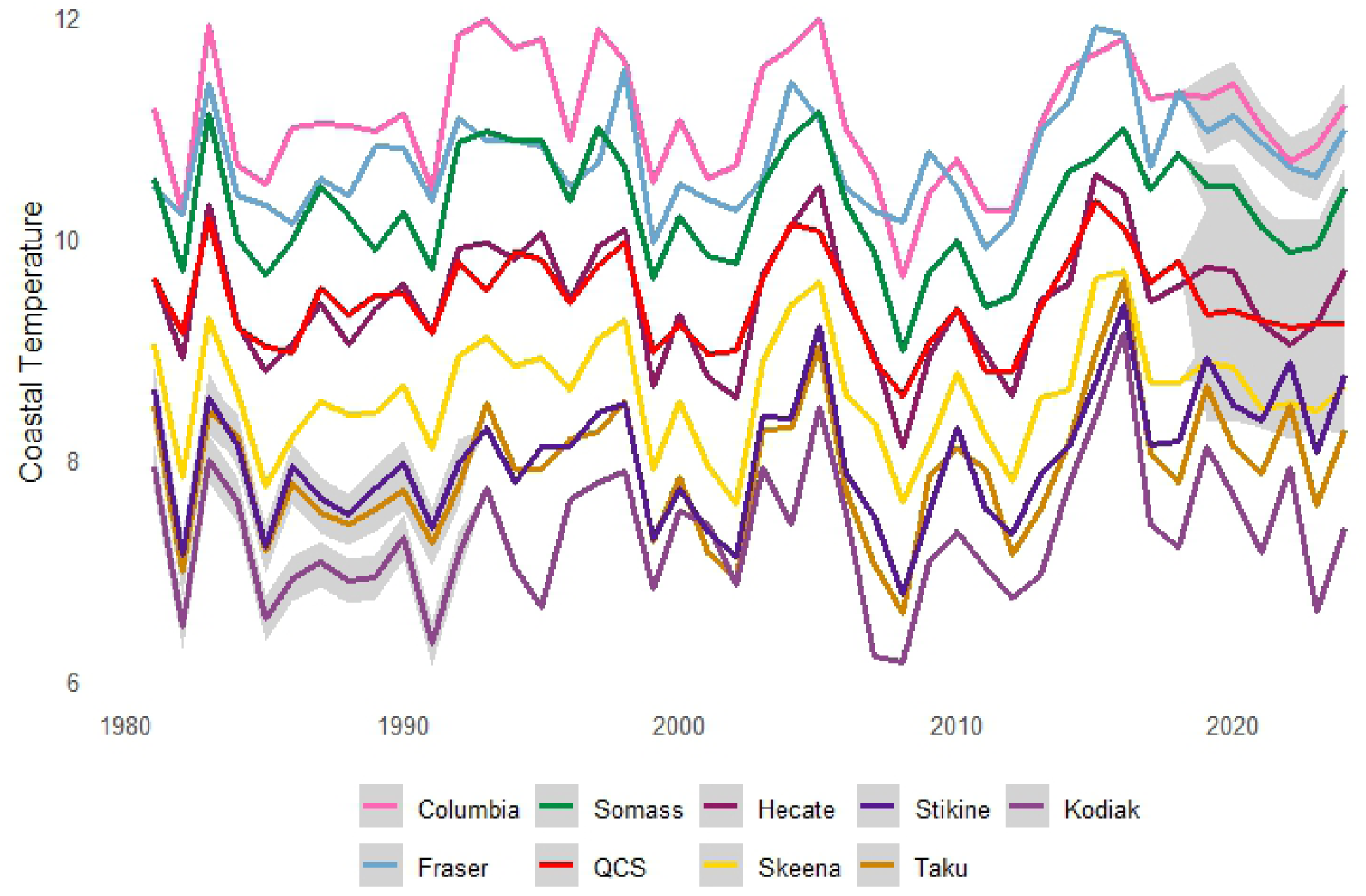
Coastal Temperatures by region in °C, colours indicate the region. Imputed data shown with 50% credible intervals.

**Figure S10:**
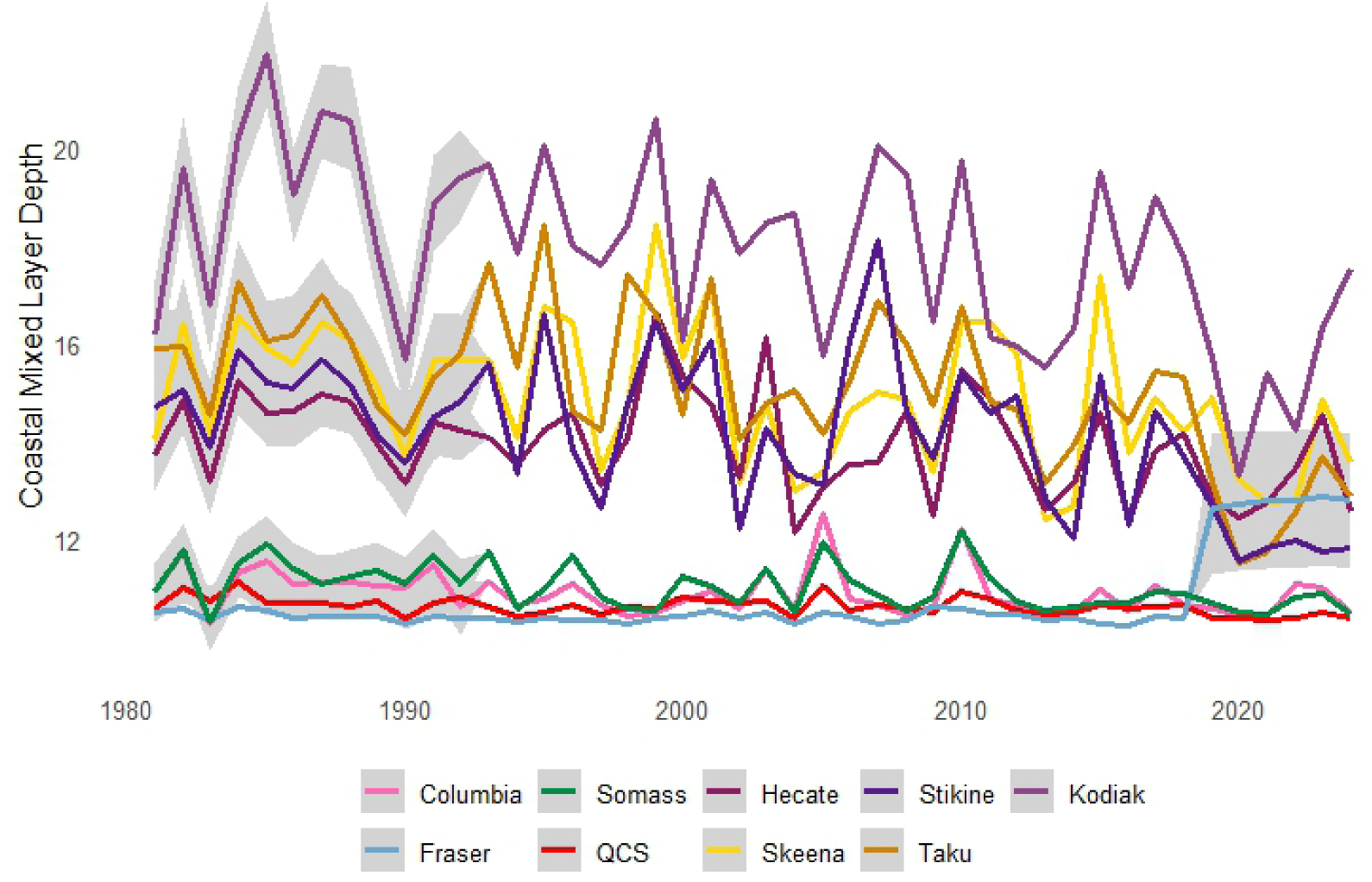
Coastal Mixed Layer Depth by region in meters, colours indicate the region. Imputed data shown with 50% credible intervals.

**Figure S11:**
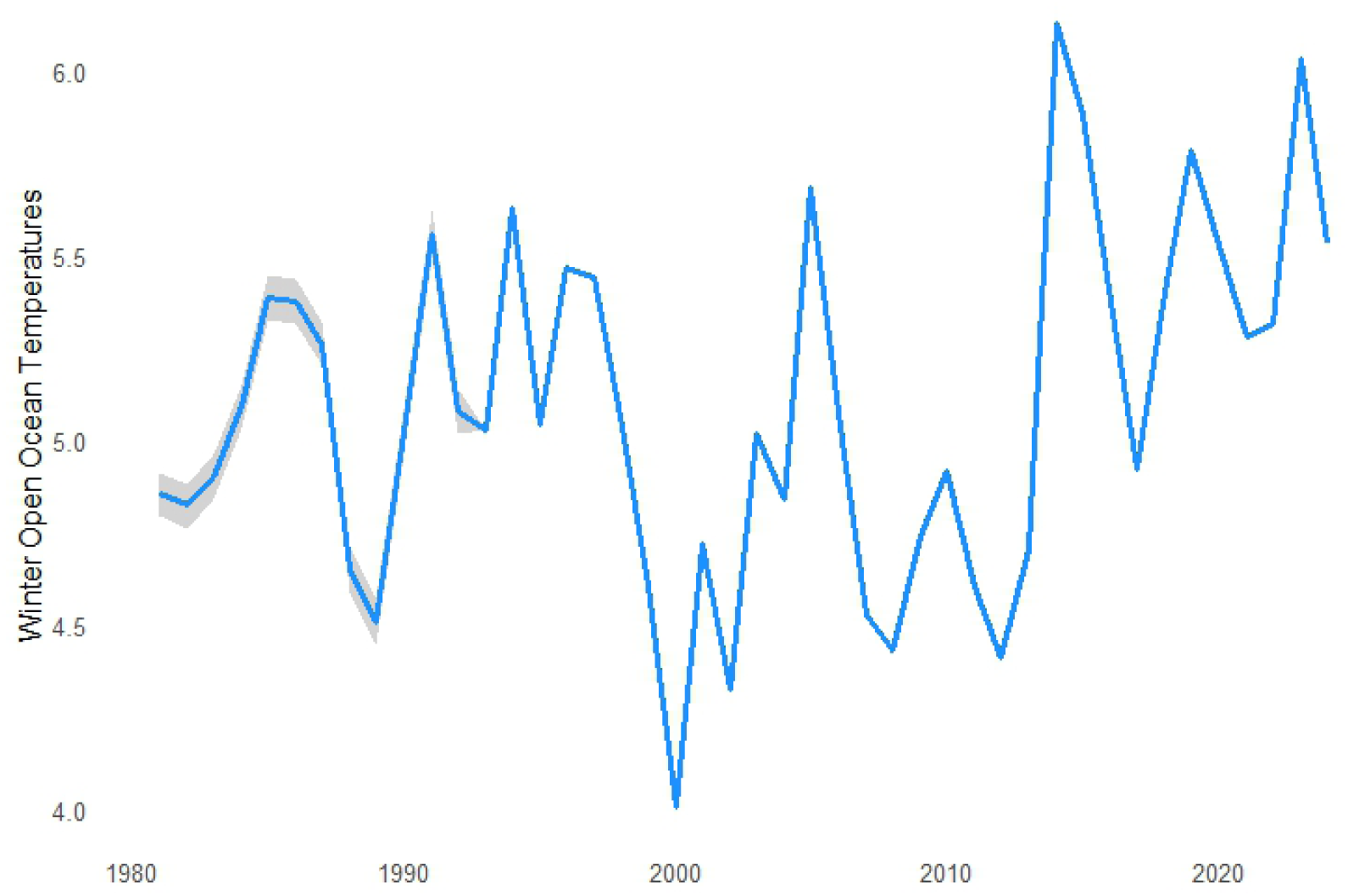
Winter Open Ocean Temperatures in °C, imputed data shown with 50% credible interval.

**Figure S12:**
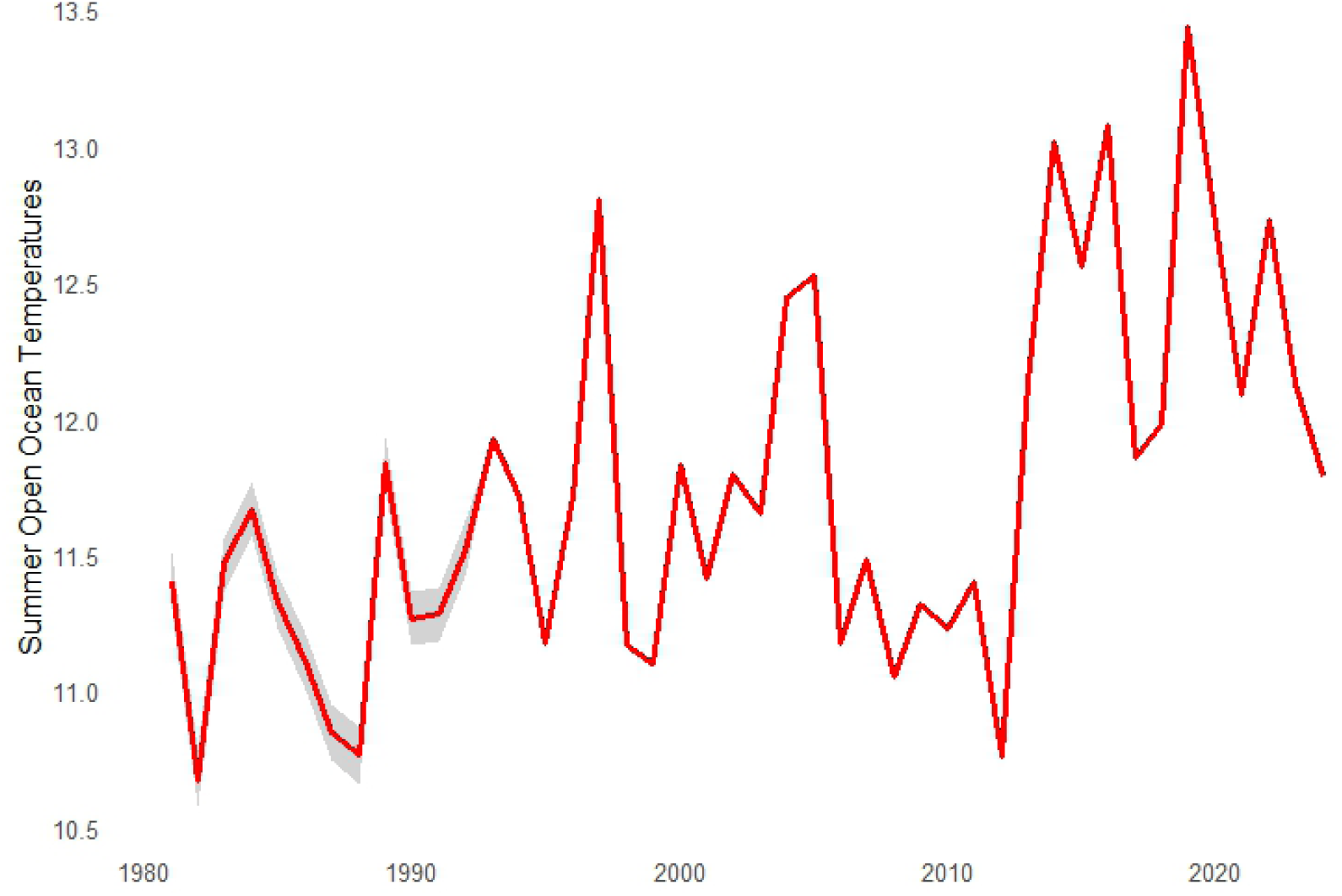
Summer Open Ocean Temperatures in °C, imputed data shown with 50% credible interval.

**Figure S13:**
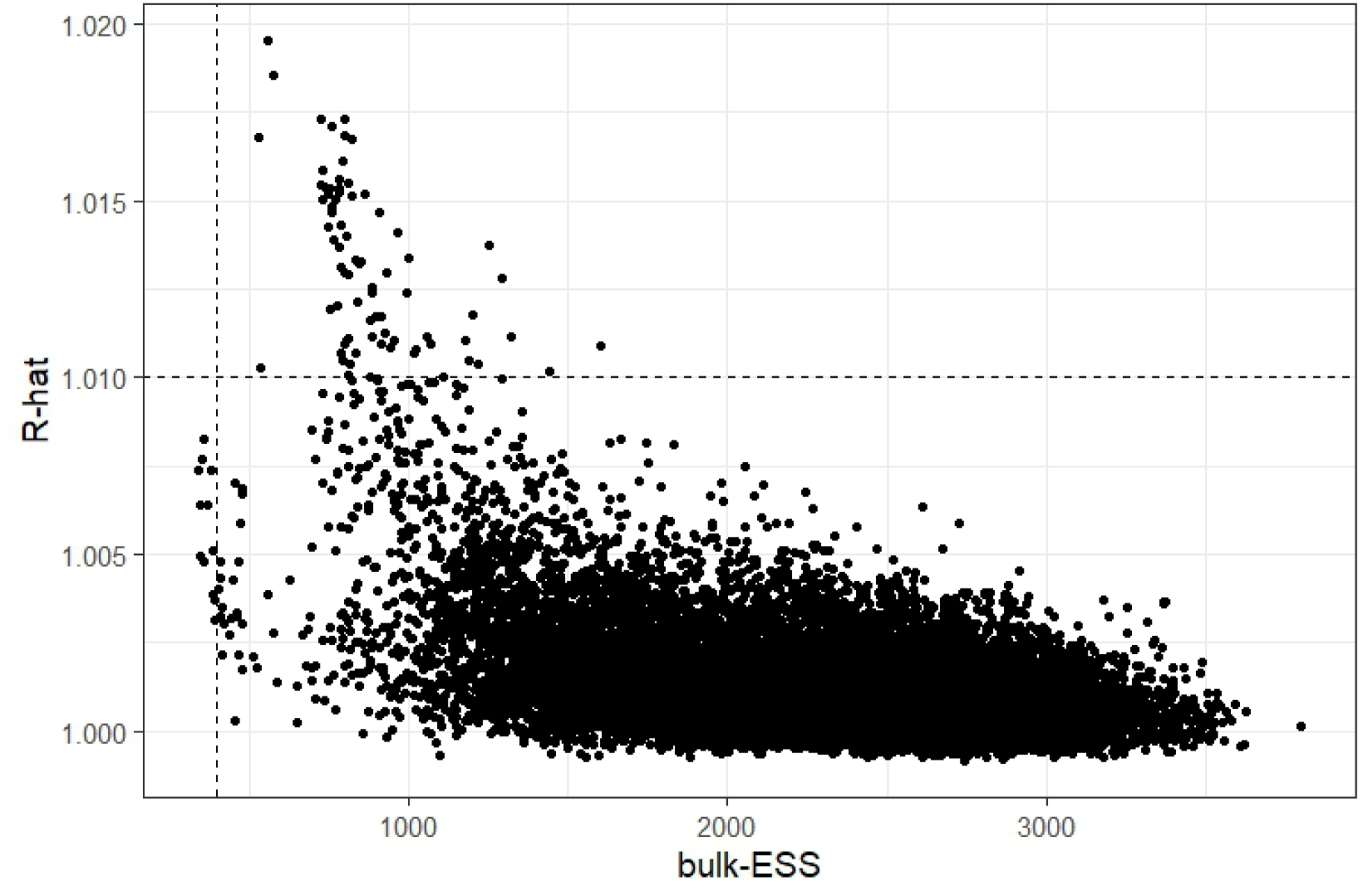
R-hat and the bulk Effective Sample Size for the model fit. The majority of draws is below a R-hat of 1.01 and a bulk ESS indicated by the dotted lines.

**Figure S14:**
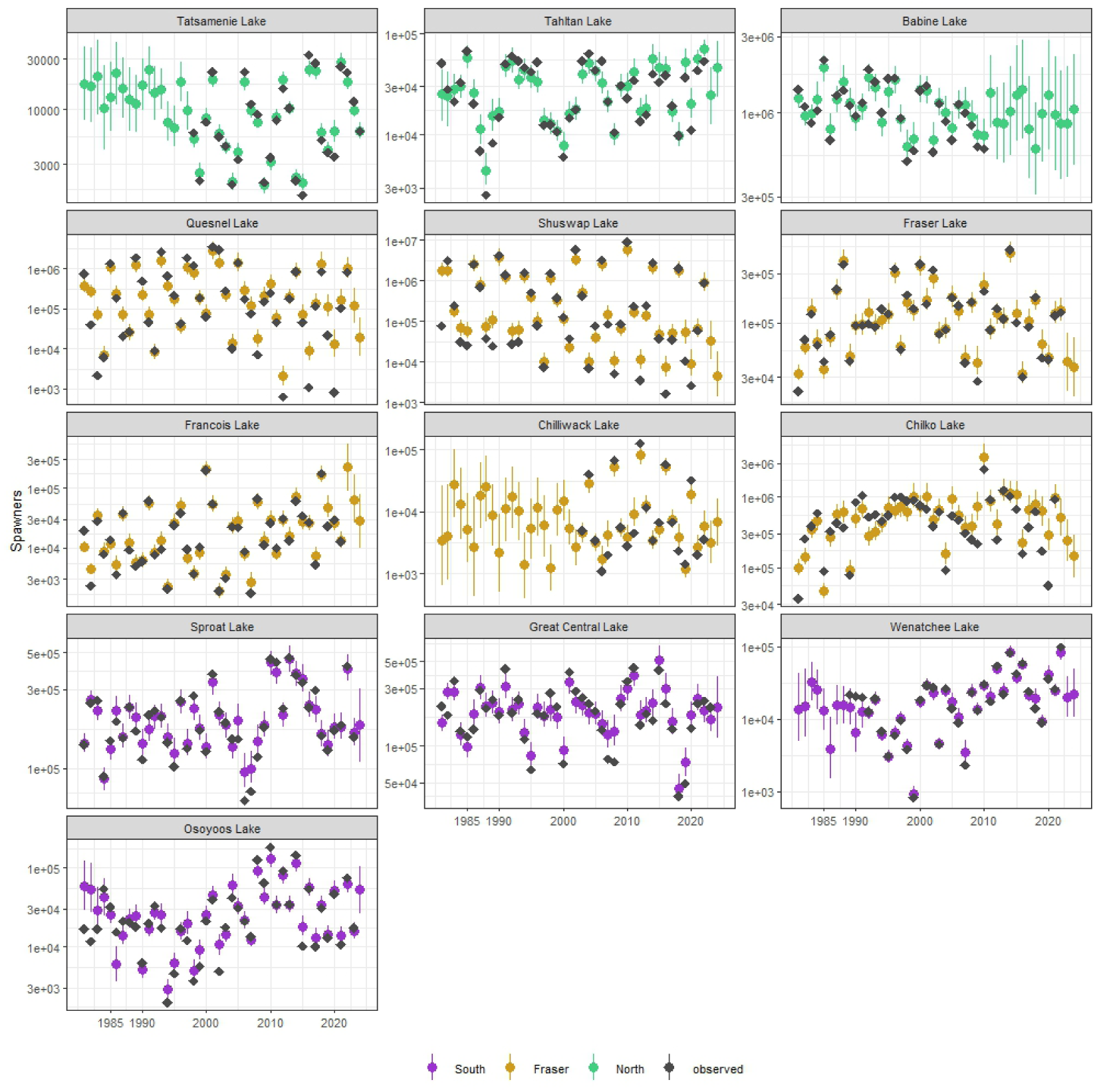
Model estimates of spawners vs. observed data in grey, colour coding by domains, Southern (purple), Fraser (gold) and Northern (green).

**Figure S15:**
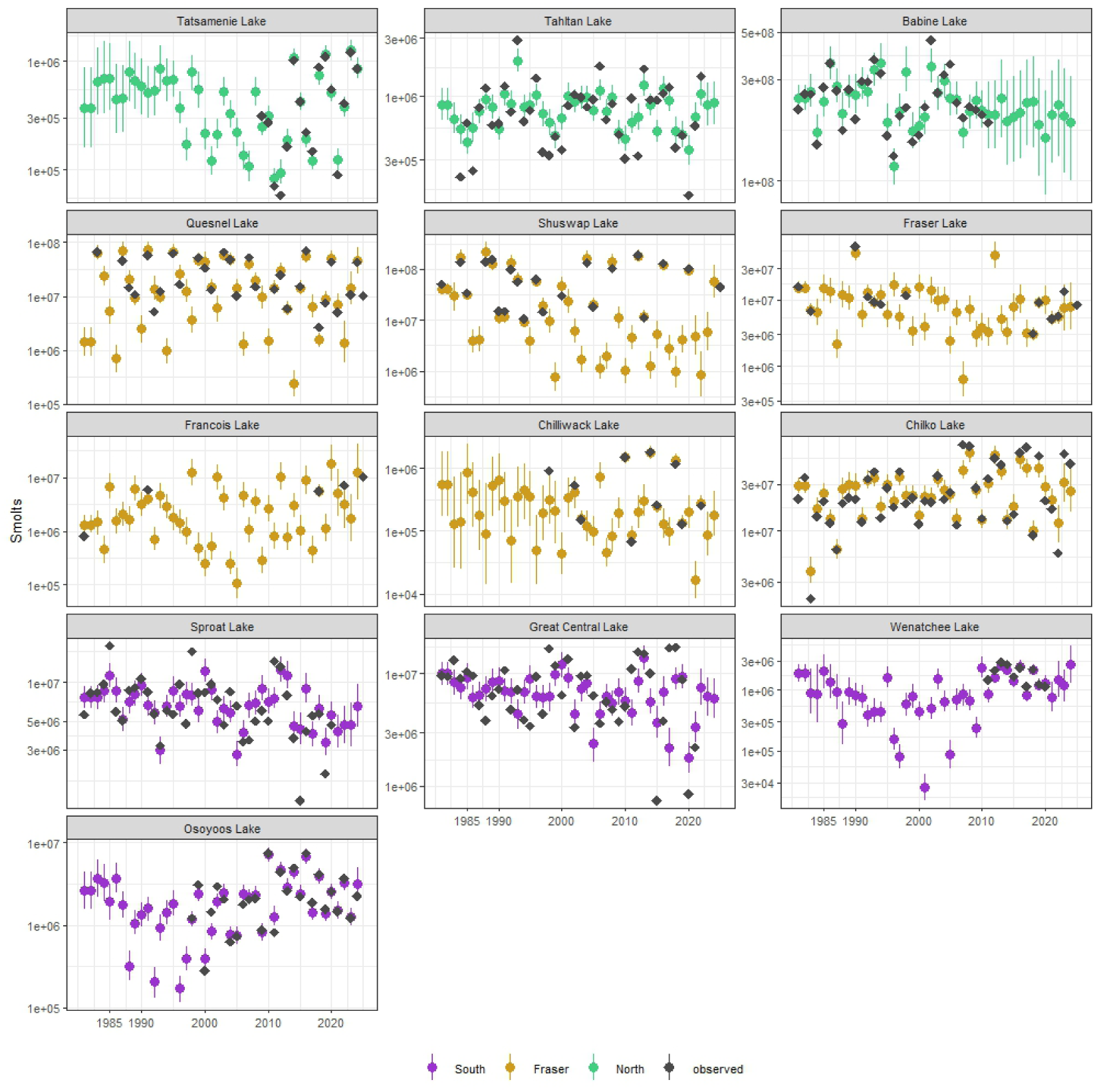
Model estimates of smolts vs. observed data in grey, colour coding by domains, Southern (purple), Fraser (gold) and Northern (green).

**Figure S16:**
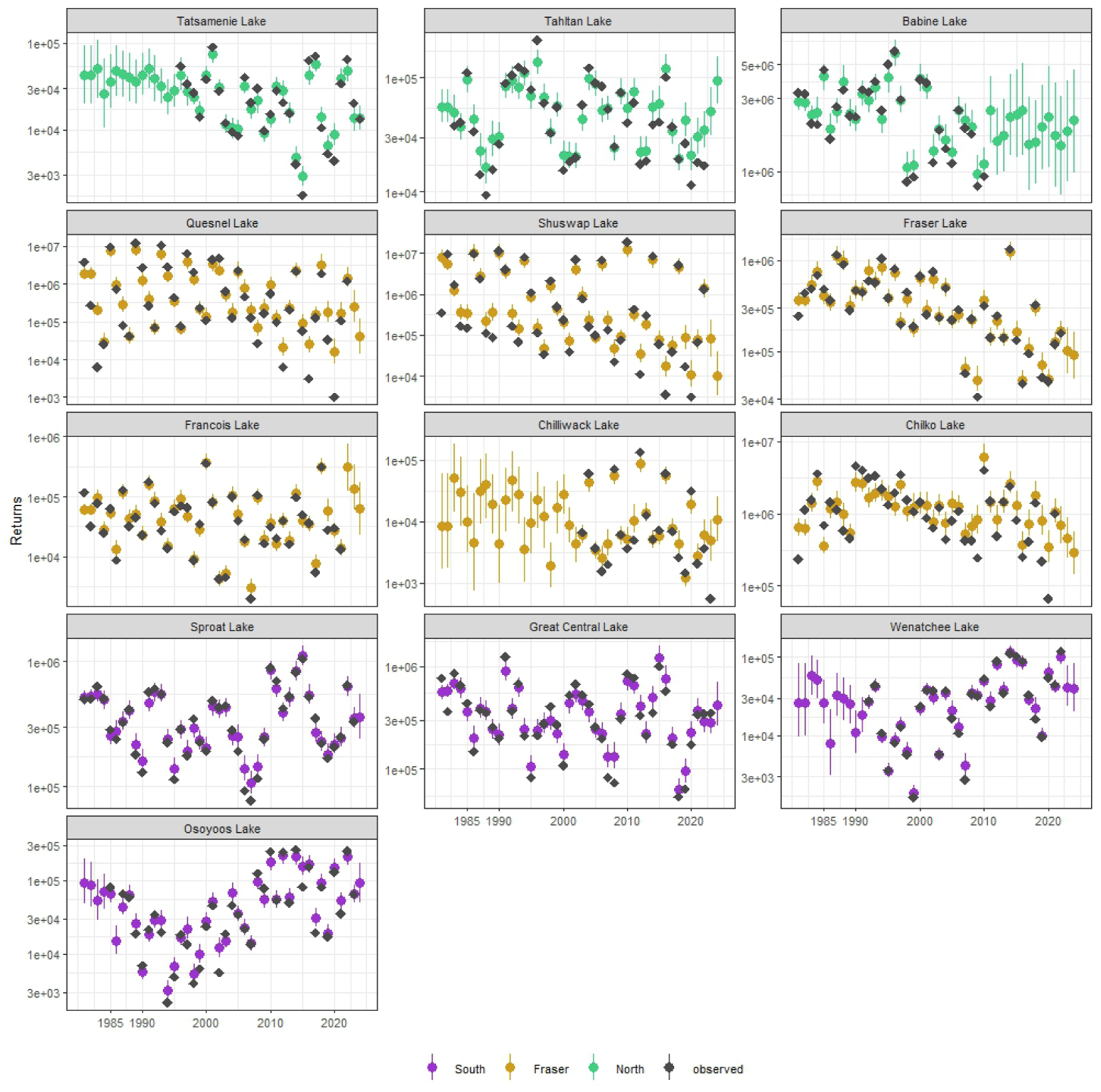
Model estimates of returns vs. observed data in grey, colour coding by domains, Southern (purple), Fraser (gold) and Northern (green).

### 5.3 Priors

**Table S1:** Priors used for imputing missing environmental data, and the return migration temperature to discharge link. The second section are the priors for imputing missing harvest/fishing rates.

| Category | Parameter | Prior |
| --- | --- | --- |
| Environmental imputation | SST_latent | $N(0,1)$ |
| Environmental imputation | TEMP_latent_a | $N(0,0.1)$ |
| Environmental imputation | TEMP_SST_b | $N(1,0.5)$ |
| Environmental imputation | TEMP_latent_sigma | $\text{LogN}(\log(0.5), 0.4)$ |
| Environmental imputation | MLDbccm_latent | $N(0,1)$ |
| Environmental imputation | MLD_latent_a | $N(0,0.1)$ |
| Environmental imputation | MLD_MLDbccm_b | $N(1,0.5)$ |
| Environmental imputation | MLD_TEMP_b | $N(-1,0.5)$ |
| Environmental imputation | MLD_latent_sigma | $\text{LogN}(\log(0.5), 0.4)$ |
| Environmental imputation | TEMPwoo_latent_a | $N(0,0.1)$ |
| Environmental imputation | TEMPwoo_SSTwoo_b | $N(1,0.5)$ |
| Environmental imputation | TEMPwoo_latent_sigma | $\text{Exp}(1)$ |
| Environmental imputation | TEMPsoo_latent_a | $N(0,0.1)$ |
| Environmental imputation | TEMPsoo_SSTsoo_b | $N(1,0.5)$ |
| Environmental imputation | TEMPsoo_latent_sigma | $\text{Exp}(1)$ |
| Environmental imputation | FTwre_latent_a | $N(0,0.1)$ |
| Environmental imputation | FTwre_GCMTwre_b | $N(1,0.5)$ |
| Environmental imputation | FTwre_latent_sigma | $\text{Exp}(1)$ |
| Environmental imputation | FTsre_latent_a | $N(0,0.1)$ |
| Environmental imputation | FTsre_GCMTsre_b | $N(1,0.5)$ |
| Environmental imputation | FTsre_latent_sigma | $\text{Exp}(1)$ |
| Environmental imputation | FTum_latent_a | $N(0,0.1)$ |
| Environmental imputation | FTum_GCMTum_b | $N(1,0.5)$ |
| Environmental imputation | FTum_latent_sigma | $\text{Exp}(1)$ |
| Environmental imputation | FDum_latent_a | $N(0,0.1)$ |
| Environmental imputation | FDum_GCMDum_b | $N(1,0.5)$ |
| Environmental imputation | FDum_latent_sigma | $\text{Exp}(1)$ |
| Environmental link | FTum_FDum_latent_a | $N(0,0.1)$ |
| Environmental link | FTum_FDum_b | $N(-0.5, 0.5)$ |
| Environmental link | FTum_FDum_latent_sigma | $\text{Exp}(\log(0.5), 0.4)$ |
| Fishing imputation | L_corr_fishing | $\text{LKJ\_Cholesky}(2)$ |
| Fishing imputation | sigma_fishing | $\text{Exp}(1)$ |
| Fishing imputation | fishing_logit_mu | $N(0,1)$ |
| Fishing imputation | fishing_prop_logit_latent | $N(0,1)$ |

**Table S2:** Environmental parameter priors used in the model.

| Category | Parameter | Prior |
| --- | --- | --- |
| Environmental effects | b_FTwre_hyper_mu | $N(0, 0.5)$ |
| Environmental effects | b_FTwre_hyper_sigma | $\text{LogN}(\log(0.25), 0.2)$ |
| Environmental effects | b_FTwre_hyper_z | $N(0, 1)$ |
| Environmental effects | b_FTwre_sigma | $\text{LogN}(\log(0.5), 0.2)$ |
| Environmental effects | b_FTwre_z | $N(0, 1)$ |
| Environmental effects | b_FTsre_hyper_mu | $N(0, 0.5)$ |
| Environmental effects | b_FTsre_hyper_sigma | $\text{LogN}(\log(0.25), 0.2)$ |
| Environmental effects | b_FTsre_hyper_z | $N(0, 1)$ |
| Environmental effects | b_FTsre_sigma | $\text{LogN}(\log(0.5), 0.2)$ |
| Environmental effects | b_FTsre_z | $N(0, 1)$ |
| Environmental effects | b_TEMPcs_hyper_mu | $N(0, 0.5)$ |
| Environmental effects | b_TEMPcs_hyper_sigma | $\text{LogN}(\log(0.5), 0.2)$ |
| Environmental effects | b_TEMPcs_hyper_z | $N(0, 1)$ |
| Environmental effects | b_TEMPcs_sigma | $\text{LogN}(\log(0.5), 0.2)$ |
| Environmental effects | b_TEMPcs_z | $N(0, 1)$ |
| Environmental effects | b_MLDcs_hyper_mu | $N(0, 0.5)$ |
| Environmental effects | b_MLDcs_hyper_sigma | $\text{LogN}(\log(0.5), 0.2)$ |
| Environmental effects | b_MLDcs_hyper_z | $N(0, 1)$ |
| Environmental effects | b_MLDcs_sigma | $\text{LogN}(\log(0.5), 0.2)$ |
| Environmental effects | b_MLDcs_z | $N(0, 1)$ |
| Environmental effects | b_TEMPwoo_hyper_mu | $N(0, 0.5)$ |
| Environmental effects | b_TEMPwoo_hyper_sigma | $\text{LogN}(\log(0.5), 0.2)$ |
| Environmental effects | b_TEMPwoo_hyper_z | $N(0, 1)$ |
| Environmental effects | b_TEMPwoo_sigma | $\text{LogN}(\log(0.5), 0.2)$ |
| Environmental effects | b_TEMPwoo_z | $N(0, 1)$ |
| Environmental effects | b_TEMPsoo_hyper_mu | $N(0, 0.5)$ |
| Environmental effects | b_TEMPsoo_hyper_sigma | $\text{LogN}(\log(0.5), 0.2)$ |
| Environmental effects | b_TEMPsoo_hyper_z | $N(0, 1)$ |
| Environmental effects | b_TEMPsoo_sigma | $\text{LogN}(\log(0.5), 0.2)$ |
| Environmental effects | b_TEMPsoo_z | $N(0, 1)$ |
| Environmental effects | b_FTum_hyper_mu | $N(0, 0.5)$ |
| Environmental effects | b_FTum_hyper_sigma | $\text{LogN}(\log(0.5), 0.2)$ |
| Environmental effects | b_FTum_hyper_z | $N(0, 1)$ |
| Environmental effects | b_FTum_sigma | $\text{LogN}(\log(0.5), 0.2)$ |
| Environmental effects | b_FTum_z | $N(0, 1)$ |
| Environmental effects | b_FDum_hyper_mu | $N(0, 0.5)$ |
| Environmental effects | b_FDum_hyper_sigma | $\text{LogN}(\log(0.5), 0.2)$ |
| Environmental effects | b_FDum_hyper_z | $N(0, 1)$ |
| Environmental effects | b_FDum_sigma | $\text{LogN}(\log(0.5), 0.2)$ |
| Environmental effects | b_FDum_z | $N(0, 1)$ |

**Table S3:**
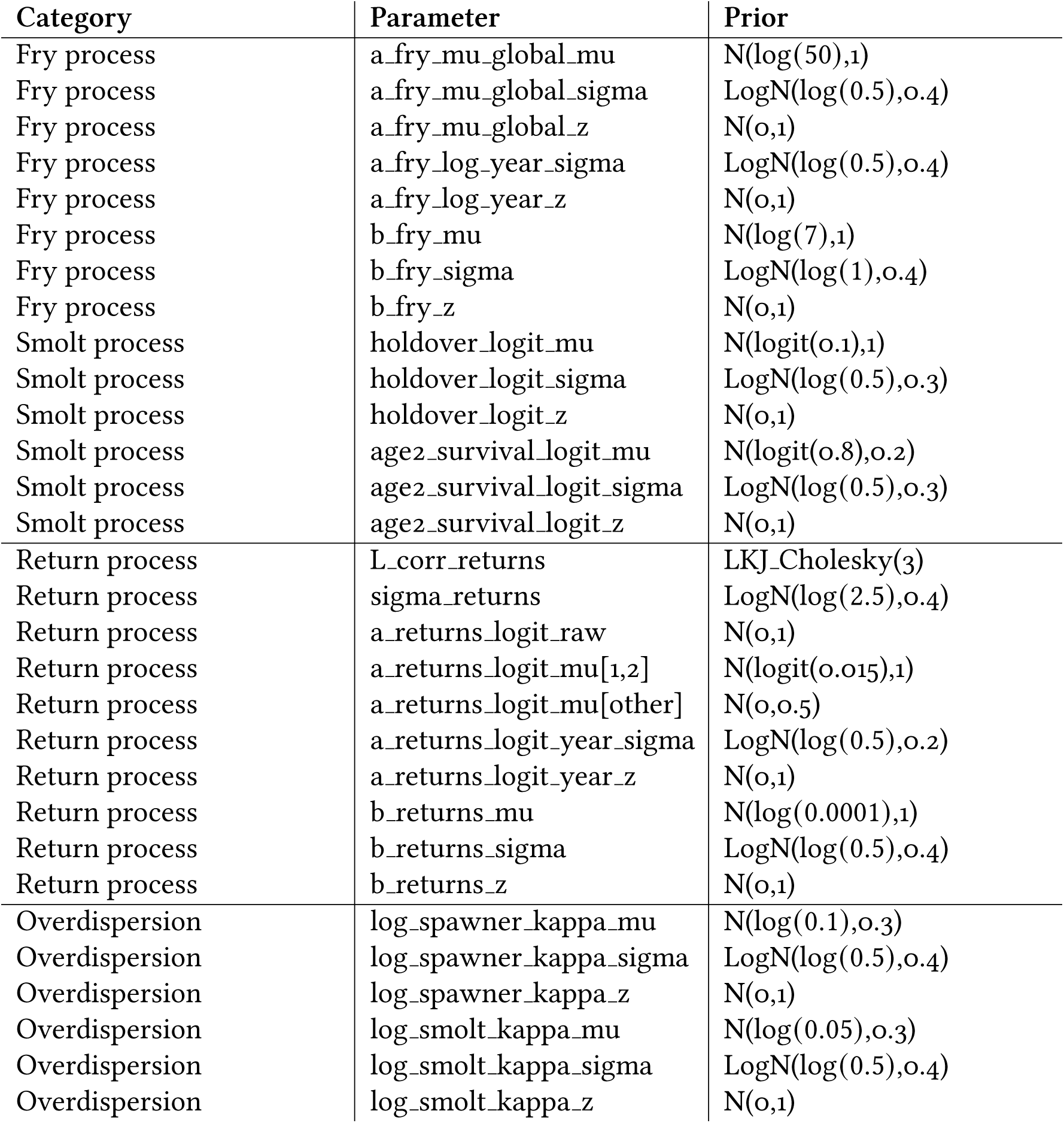
Process model parameter priors for Spawners to Fry, Fry to Smolts, and Smolts to Returns. The third section lists the overdispersions for the observation model.

### 5.4 Simulation Model Supplement

Because the true environmental effects influencing productivity are unknown in our empirical dataset, we tested model performance using simulated data for which the covariate effects were specified *a priori*. This simulation-recovery exercise allowed us to assess whether the model could accurately estimate the magnitude and direction of environmental effects.

Simulated datasets of spawner abundance, smolt abundance, and age-class structure for all populations were generated using the same life-cycle process model structure used in the analysis, but implemented in R (R Core Team 2025). Environmental covariate time series were taken from those used in the fitted model, with missing values replaced by the corresponding imputed values from the Stan model. All parameters other than the environmental-effect coefficients were fixed at their posterior median estimates from the fitted model. This included all year-level random effects. This was done to generate population dynamics that broadly matched the magnitude and variability of the populations in our study, while allowing the environmental effects on productivity to be specified directly.

Covariate parameters for each population were drawn from normal distributions centred on specified means to allow variability among populations. Parameters were chosen to span a range of positive, negative and null responses that differed from those estimated in the Stan model. Mean effects were set to 0 for winter freshwater temperature, -0.5 for summer freshwater temperature, -0.1 for coastal temperature, -0.5 for coastal mixed layer depth, 0.2 for winter open-ocean temperature, -0.2 for summer open ocean temperature, -0.25 for return-migration temperature, and 0.1 for return migration discharge. All coefficients shared a common standard deviation (0.05), with the exception of winter freshwater temperature, for which coefficients were fixed at 0. This allowed variation among populations, while ensuring that one covariate had no effect, allowing us to test whether the model could identify null relationships. Because our objective was to evaluate recovery of environmental-effect coefficients rather than to estimate the hierarchical structure, we omitted the domain-level hierarchical structure when setting the environmental effects in our simulation.

Simulations were initialized with initial abundances estimated in the empirical model and we included a 20-year burn-in period to reduce the influence of these initial conditions. During the burn-in period, annual environmental conditions were sampled randomly from the observed covariate time series. This was followed by a 44-year simulation using the environmental covariate time series used in the empirical model (with imputed data as described above). The simulated datasets were then analyzed using a simplified version of the Stan model that omitted missing-data imputation because complete covariate time series were available for all simulated observations. Posterior distributions were obtained for the environmental-effect parameters and compared with the values used to generate the simulated data.

The model generally recovered the environmental-effect parameters used to generate the simulated data. Across all populations and covariates, the true parameter value was within the estimated 95% credible interval 97 of 104 times (93.3%; Figure S17). Parameters that were not recovered were spread across covariates and populations, suggesting no systematic bias in parameter estimation. Notably, all winter freshwater temperature effects were recovered, despite this covariate being assigned a true effect of zero in the simulation. This indicates that the model was able to identify null environmental effects.

**Figure S17:**
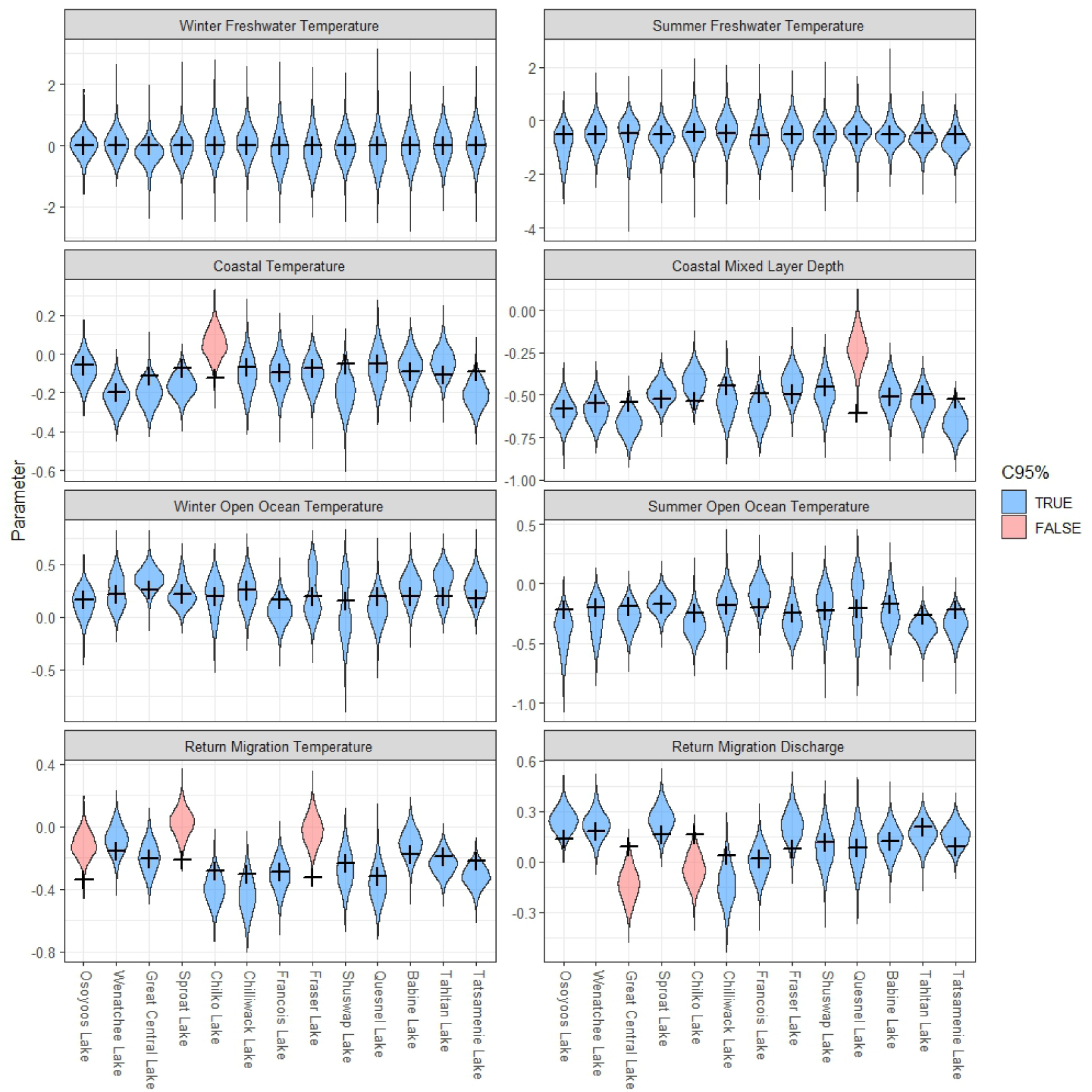
Posterior estimate distributions of parameters from the simulated populations dynamics using the lifecycle model. Black crosses indicate the parameter value set in the data simulation. Blue designates that the set value was within the 95% credible interval of the posterior estimates for a given population and parameter, red that it was not.

### 5.5 Shapley Values

To assess the contributions of the combined environmental effects, spawner abundance effects and random effects to the model estimates of returns, we estimated expected total returns under a series of hypothetical scenarios that included all combinations of these components (Table S4). Each component was either set at a reference value (median population specific spawners; median population specific environmental conditions; random year effects set to 0) or to its year-specific value. These hypothetical scenarios used the estimated latent spawner abundances as predictors, rather than the observed values, to remove observation error and to avoid missing observations. Shapley values (Shapley 1952) were then calculated from the differences in expected returns across these counterfactual scenarios. Because the Shapley values are based on expected returns rather than observed returns, they do not account for observation error. However, the comparisons of observed vs. expected returns are shown in Figure S16.

**Table S4:** Hypothetical scenarios used to compute Shapley contributions to predicted returns. 0 indicates reference value, 1 indicates the year-specific values.

| Scenario | Spawners (*S*) | Environment (*E*) | Random effects (*R*) |
| --- | --- | --- | --- |
| Reference | 0 | 0 | 0 |
| *S* | 1 | 0 | 0 |
| *E* | 0 | 1 | 0 |
| *R* | 0 | 0 | 1 |
| *S + E* | 1 | 1 | 0 |
| *S + R* | 1 | 0 | 1 |
| *E + R* | 0 | 1 | 1 |
| *S + E + R* | 1 | 1 | 1 |

The Shapley value represents the average marginal contribution of each component, averaged across all possible scenarios in which the component is either included or excluded. Effect contributions therefore represent deviations (as numbers of returning fish) from the population specific median reference estimates. Annual Shapley values for each population were subsequently used to quantify interannual variance and to compute the proportional contribution of each component to the total interannual return variability.

## Author Statement

J.F.F.: Conceptualization, Project administration, Data curation, Methodology, Investigation, Formal analysis, Code, Validation, Visualization, Writing original draft, Editing T.T.: Conceptualization, Project administration, Data curation, Methodology, Investigation, Formal analysis, Code, Validation, Visualization, Writing original draft, Editing C.F.: Conceptualization, Funding acquisition, Project administration, Supervision, Resources, Data curation, Methodology, Investigation, Formal analysis, Code, Validation, Visualization, Writing original draft, Editing B.C.: Conceptualization, Funding acquisition, Project administration, Supervision, Methodology, Visualization, Writing original draft, Editing A.M.H.: Conceptualization, Funding acquisition, Data curation, Methodology, Validation, Editing G.L.O.: Conceptualization, Resources, Methodology, Editing D.T.S.: Conceptualization, Resources, Editing H.W.S.: Conceptualization, Resources, Editing P.L.T.: Conceptualization, Funding acquisition, Project administration, Supervision, Methodology, Investigation, Formal analysis, Code, Validation, Visualization, Writing original draft, Editing

## Acknowledgements

We acknowledge the late Dr. Kim Hyatt, who originally identified the benefit of integrating data on stage-specific survival across a network of sockeye salmon populations. His scientific curiosity lay the groundwork for the analysis we pursued here. We are grateful to the biologists and field technicians who have collected salmon data over the previous decades. Adam Brennan, Nick Brown, Jason Calvert, Marissa Glavas, Jodie Mackenzie-Grieve, Helen Liu, Athena Ogden, Lucas Pon, Eric Taylor, Catherine Willard, and the Pacific Salmon Commission Secretariat provided juvenile and adult abundance data. We are also grateful to the Okanagan Nation Alliance and COBTWG for providing data and insight. Skip McKinnell provided helpful insight on marine

environmental variables. Angelica Peña provided down-scaled marine environmental data, Markus Schnorbus provided down-scaled freshwater environmental data from the Pacific Climate Impacts Consortium. Josie Iacarella helped guide initial project development.

## Data Availability Statement

Data, and Stan and R analysis code used in this project are available at https://doi.org/10.5281/zenodo.21795158 and https://github.com/SiRE-P/PSSIsalmonclimatepublic.git. Fisheries data for the Osoyoos Lake population are owned by the Okanagan Nation Alliance. Please contact Karilyn Alex to request access to the data.

